# PI3Kβ inhibition restores ALK inhibitor sensitivity in ALK-rearranged lung cancer

**DOI:** 10.1101/2021.03.18.435801

**Authors:** Sarang S. Talwelkar, Mikko I. Mäyränpää, Julia Schüler, Nora Linnavirta, Annabrita Hemmes, Simone Adinolfi, Matti Kankainen, Wolfgang Sommergruber, Anna-Liisa Levonen, Jari Räsänen, Aija Knuuttila, Emmy W. Verschuren, Krister Wennerberg

## Abstract

For non-small cell lung cancer (NSCLC) patients with *ALK*-rearranged tumors, treatment with ALK inhibitors can improve outcomes. However, clinical resistance typically develops over time, and in the majority of cases resistance mechanisms are ALK-independent. We generated tumor cell cultures from multiple regions of an *ALK*-rearranged clinical tumor specimen, and deployed functional drug screens to identify modulators of resistance to ALK inhibitors. This identified a role for PI3Kβ and EGFR in regulating resistance to ALK inhibition. Furthermore, inhibition of ALK elicited activation of EGFR, and inhibition of PI3Kβ rescued EGFR-mediated ALK inhibitor resistance. In *ALK*-rearranged primary cultures, cell lines and *in vivo* xenograft models, combined inhibition of ALK and PI3Kβ prevented compensatory MAPK and PI3K-AKT pathway reactivation and selectively targeted the cancer cells. The combinatorial effect was seen even in the background of *TP53* mutations and in epithelial-mesenchymal transformed cells. In conclusion, combinatorial ALK and PI3Kβ inhibitor treatment carries promise as a treatment for *ALK*-rearranged NSCLC.

## INTRODUCTION

In 3-7% of NSCLCs, *ALK* gene rearrangements lead to the expression of oncogenic fusion proteins that confer constitutive activity of the ALK kinase domain ^1^. Aberrant ALK activity increases the MAPK and PI3K-AKT oncogenic signaling pathways ^2, 3^. Treatment of *ALK*-rearranged NSCLC with the first-generation ALK inhibitor crizotinib associates with favorable response in around 70% of the patients, although resistance and clinical relapse typically occurs within a year^4^. Moreover, around 5% of *ALK*-rearranged NSCLC patients show progressive disease following crizotinib treatment ^4^. Mechanisms for intrinsic resistance are poorly understood, but have been proposed to relate to e.g. the precise *ALK* fusion variant^5^, or *TP53* mutations which frequently co-occur with *ALK* rearrangements^6–8^. Among tumors that acquire resistance to crizotinib, about 30% exhibit *ALK*-dependent resistance mechanisms, particularly *ALK* mutations and/or amplification ^9, 10^. In the remainder, resistance is driven by diverse mechanisms, including activation of bypass signaling pathways (e.g., EGFR, KIT, IGF-1R, and GPCRs), epithelial-to-mesenchymal transition (EMT), a histopathology switch, or increased activity of drug efflux pumps ^2^.

For *ALK*-rearranged NSCLCs exhibiting ALK-dependent acquired resistance, second- or third-generation ALK inhibitors that retain activity in the presence of resistance mutations to first-line inhibitors offer optional “sequential” treatment opportunities. Nevertheless, eventually most patients relapse due to development of compound ALK mutations (∼40%) or loss of dependency of tumor cells on ALK (∼60%), regardless of the timing and type of the prescribed drug ^11^. However, the mechanistic drivers for resistance, such as bypass signaling pathways, are not addressed by clinical diagnostics, and these mechanisms are likely different between samples. To address this clinical problem, patient-derived cultures from ALK inhibitor-resistant tumors have been used to identify treatment-adaptive resistance mechanisms and combinatorial treatment approaches ^12–14^. However, as of yet, no combinatorial treatments to counter bypass signaling are available in the clinic.

In addition to pronounced inter-tumor variability in resistance mechanisms, *ALK*-rearranged lung tumors also exhibit intra-tumor heterogeneity in ALK inhibitor resistance ^2^. To understand how intra-tumoral heterogeneity may contribute to variable ALK-related drug sensitivity, we performed multiregional characterization of an *ALK*-rearranged lung tumor and corresponding tumor-derived cultures to identify combinatorial treatment options. Our study identified a novel role for PI3Kβ in mediating resistance to ALK inhibition. As PI3Kβ is an effector of multiple tyrosine kinase activities that can mediate ALK inhibitor resistance ^15–20^, including EGFR, co-targeting of PI3Kβ and ALK offers promise as a treatment for *ALK*-rearranged lung cancer.

## RESULTS

### Intratumor heterogeneity of an *ALK*-rearranged lung tumor

To capture the extent of intratumor heterogeneity and to identify treatments that can enhance sensitivity to ALK inhibition in *ALK*-rearranged lung cancer, we characterized tumor tissues (n=6) and cultures (n=4) established from multiple tumor regions (TRs) of a large size (9×12×9 cm) surgically resected tumor from a 55-year-old never-smoker female patient with *ALK*-rearranged lung adenocarcinoma with diffuse metastasis. Before surgery, the patient was treated with chemotherapy, which did not halt disease progression (Fig. 1A and S1A-B). Immunohistochemical characterization of six tumor regions was done for ALK and markers of lung adenocarcinoma (NKX2.1), epithelial (E-cadherin, pan-cytokeratin, and cytokeratin 18) and mesenchymal (vimentin) phenotypes, tumor vasculature (CD31), and basement membrane organization (collagen type IV). All but one tissue region showed a similar phenotype with cancer cells exhibiting positivity for both NKX2.1 and the epithelial marker E-cadherin. The exception was tumor region #5 (TR5), which exclusively showed aggressive lung cancer features, including aberrant basement membrane organization evidenced by enhanced collagen type IV staining, high tumor vasculature, and mesenchymal phenotype cancer cells lacking NKX2.1 expression (Fig. 1B and S1C). Importantly, TR5-derived cells exhibited predominant expression of ALK and vimentin formed tightly-packed clusters/colonies upon *ex vivo* culture, while cultures from other regions formed uniform epithelial monolayers (Fig. 1C-D and S1D), corroborating the parental mesenchymal tissue phenotype of the cancer cells in TR5.

**Figure 1.**
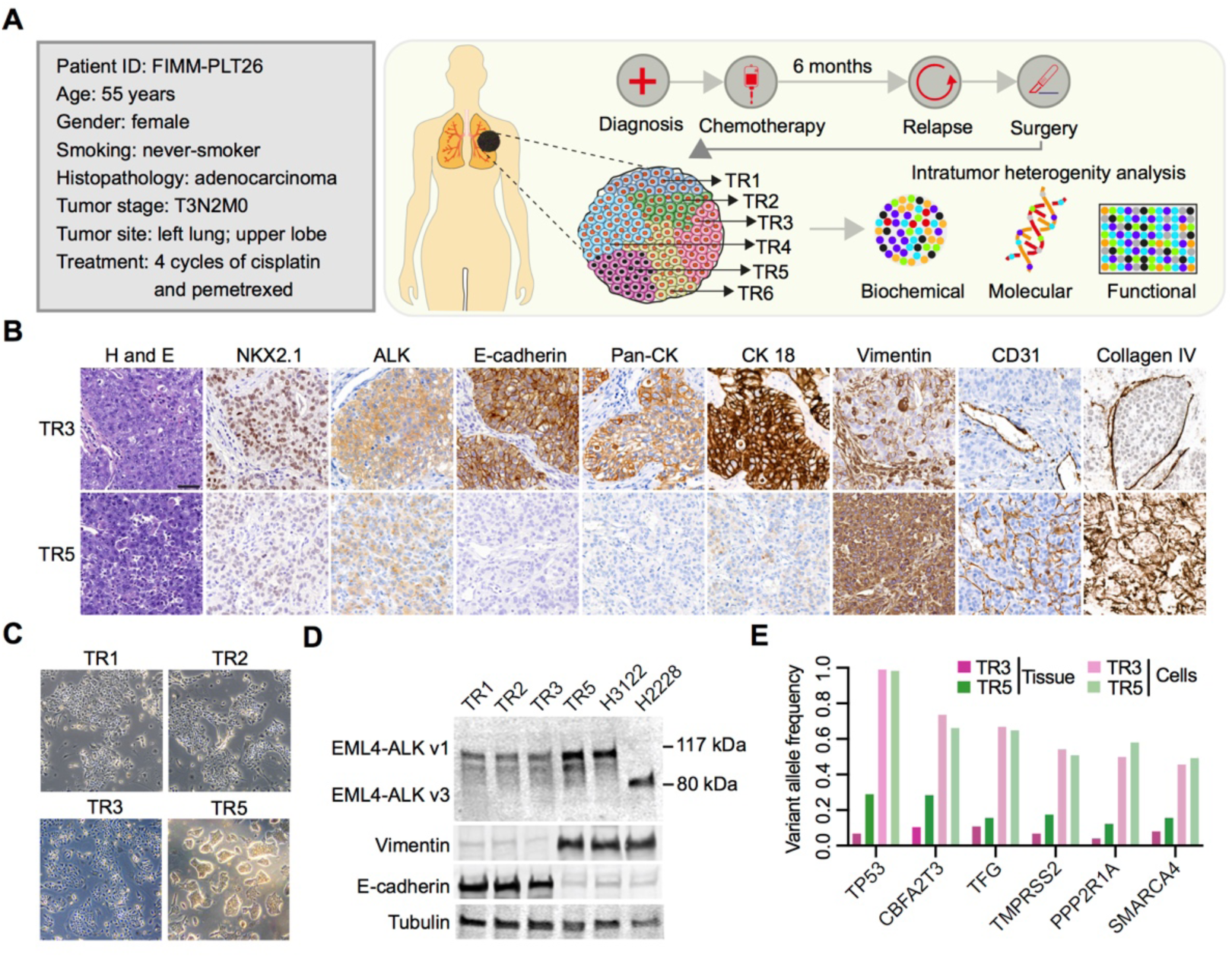
*ALK*-rearranged lung tumor analysis reveals EMT-related intratumor heterogeneity. (A) Clinical features and outline of intratumor heterogeneity analyses in a lung cancer specimen. (B) Representative images of hematoxylin and eosin (H&E) and IHC staining performed on TR3 and TR5 tissue regions. Scale bar, 50 µm. (C) Bright field pictures of cultures derived from different tissue regions. (D) Immunoblots of *ALK*-rearranged tumor-derived cultures and cell lines, probed with the indicated antibodies. Lysates from H3122 and H2228 were used as a positive control for ALK variants 1 (v1) and 3 (v3), respectively. (E) Variant allele frequencies of cancer-selective somatic mutations identified in each sample.

We next asked if TR5 cancer cells carried unique genetic aberrations to explain their divergent phenotype. Targeted next-generation sequencing of the TR3 and TR5 tumor tissues and their corresponding cultures, however, showed an identical set of somatic mutations (Fig. 1E), suggesting that the difference between TR5 and other regions was phenotypic rather than mutational. The aggressive nature of the cancer was underscored by multiple mutations in tumor suppressor genes, including *TP53*, *CBFA2T3, PPP2R1A,* and *SMARCA4*. As expected, variant allele frequencies were substantially higher in cultures than in their respective tissues, demonstrating cancer cell enrichment in cultures. This overall demonstrates intra-tumoral phenotypic heterogeneity and a genomic profile that underscores the aggressive nature of this *ALK*-rearranged lung cancer sample.

### Drug sensitivity profiling reveals intra-tumor functional heterogeneity

To understand how intratumor heterogeneity might impact cancer cell functions, we analyzed the drug sensitivities of tumor-derived cultures from four regions. Cells derived from patient-matched tumor and normal lung tissue as well as *ALK*-rearranged H3122 cells were screened with a customized panel of 527 anticancer compounds representing both investigational and FDA-approved compounds targeting a broad range of molecular targets, including more than 100 compounds targeting ALK or its effector pathways (Table S2). Matching with the patient’s clinical response, tumor-derived cultures showed lack of cancer cell-selective response to cisplatin and pemetrexed (Fig. S1E). As expected, only *ALK*-rearranged cancer cells but not normal lung epithelial cells showed sensitivity to ALK inhibitors (ALKi) (Fig. 2A-C and S2A). ALKi responses, while selective, were moderate, in agreement with the *TP53* mutant status of the cells. Correlation comparisons of drug sensitivity scores (DSSs) of the entire drug screen showed a high overall degree of correlation between *ALK*-rearranged cells, but comparison of ALKi responses demonstrated that TR5 cells exhibited lowest ALKi sensitivities and exhibited least degree of correlation with other tumor-matched cancer cells and H3122 cells (Fig. 2B and S2A-D). Comparison of the 20 most effective compounds identified for each cell model showed many hits unique for TR5-derived cells (55%), including selective sensitivity to HDAC inhibitors (n=4) (Fig. S2E).

**Figure 2.**
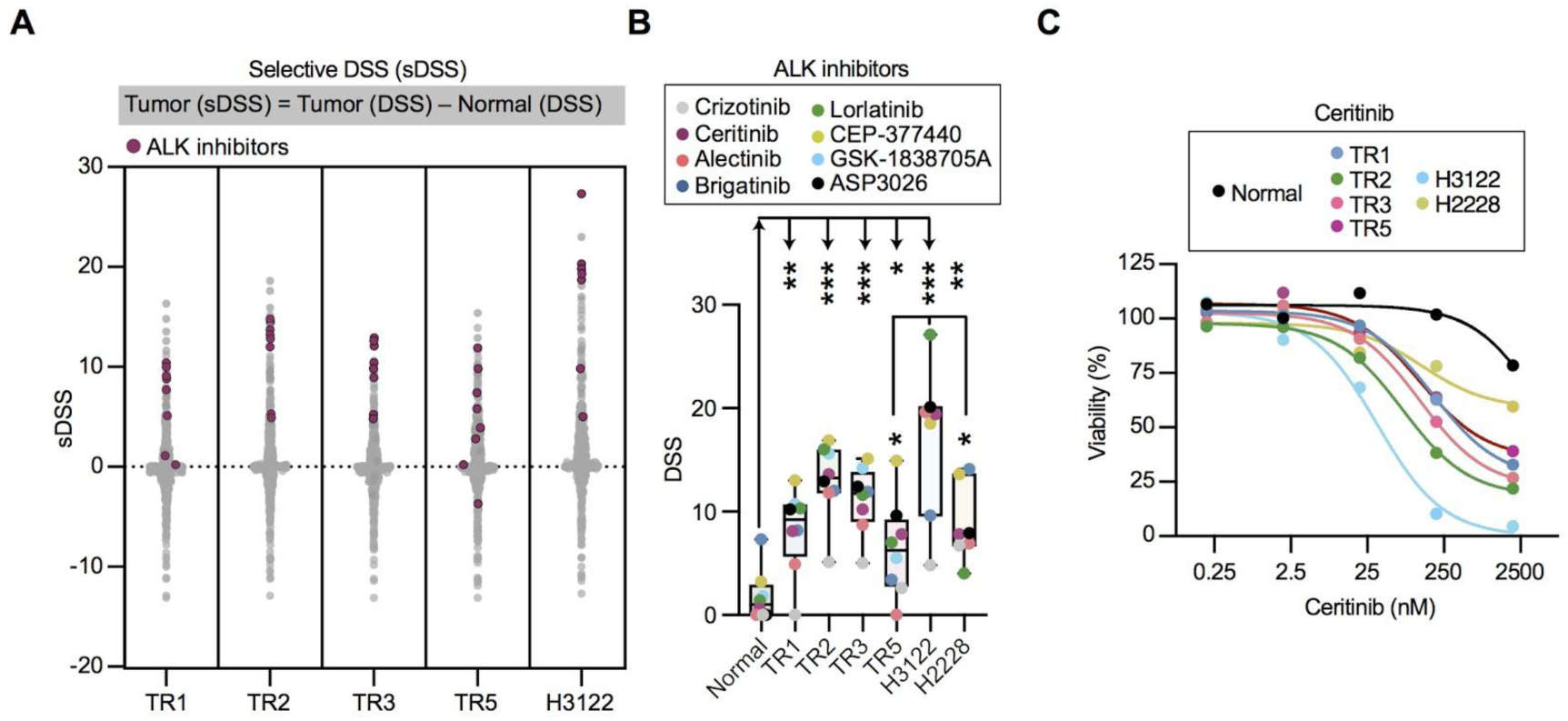
Pharmacological profiling reveals functional heterogeneity of an *ALK*-rearranged NSCLC tumor. (A) Integrated results of DSRT performed on tumor-derived and H3122 cells; cancer-selective hits were identified on the basis of selective DSSs. *ALK*-rearranged lung cancer-selective compounds targeting ALK are depicted as red. (B) The DSSs of ALKi for indicated cell types. Each dot represents an individual drug and dots are color-coded based on drug ID. GSK-1838705A was not tested for H3122 and H2228 cells. (C) Dose-response curves of normal lung, tumor-derived cells, H3122, and H2228 cells treated with the second generation ALKi, ceritinib. Whiskers represent minimum and maximum values. Student’s *t* test *p* values are * < 0.05, ** < 0.01, *** < 0.001.

In comparison to H3122, H2228 cells also showed relatively low sensitivities to ALKi, both in our study (Fig. 2B-C and S2A) as well in the GDSC dataset (Fig. S2F). Even though H2228 cells are a widely used model of *ALK*-rearranged lung cancer, causative factors underlying low ALKi sensitivities remain to our knowledge unexplored. H2228 cells carry a *TP53* mutation and a *NFE2L2* (Nrf2) G31A gain-of-function mutation, which constitutively activates the Nrf2 signaling pathway. Both mutations are associated with low sensitivity to ALKi ^7, 8, 21^. To confirm the dysregulation of the Nrf2 signaling pathway, we compared the expression of Nrf2 and its target genes at both the mRNA and protein level in H3122 and H2228 cells. This indeed showed that H2228 cells exhibited considerably higher expression of Nrf2 and Nrf2 downstream targets than H3122 cells (Fig. S2G-H). Taken together, drug sensitivity testing of tumor-derived cultures (and cell lines) exposed functional heterogeneity of this particular *ALK*-rearranged lung cancer.

### Inhibition of ALK activates EGFR in *ALK*-rearranged lung cancer cells

To assess long-term sensitivity of tumor-derived cultures to ALKi, cells were exposed to IC50 doses of five ALKi for 15 days. All tumor-derived cultures showed paradoxical cell growth as evidenced by increased colony formation in presence of ALKi (Fig. 3A-B). To understand the molecular mechanism for this limited ALKi sensitivity of tumor-derived cultures, we analyzed adaptive signaling following ceritinib treatment. Ceritinib was chosen as the most frequently used ALKi in clinical trials that investigate the efficacy of first-line combinatorial treatment of *ALK*-rearranged lung cancer, and the only second generation approved ALKi at the start of our study.

**Figure 3.**
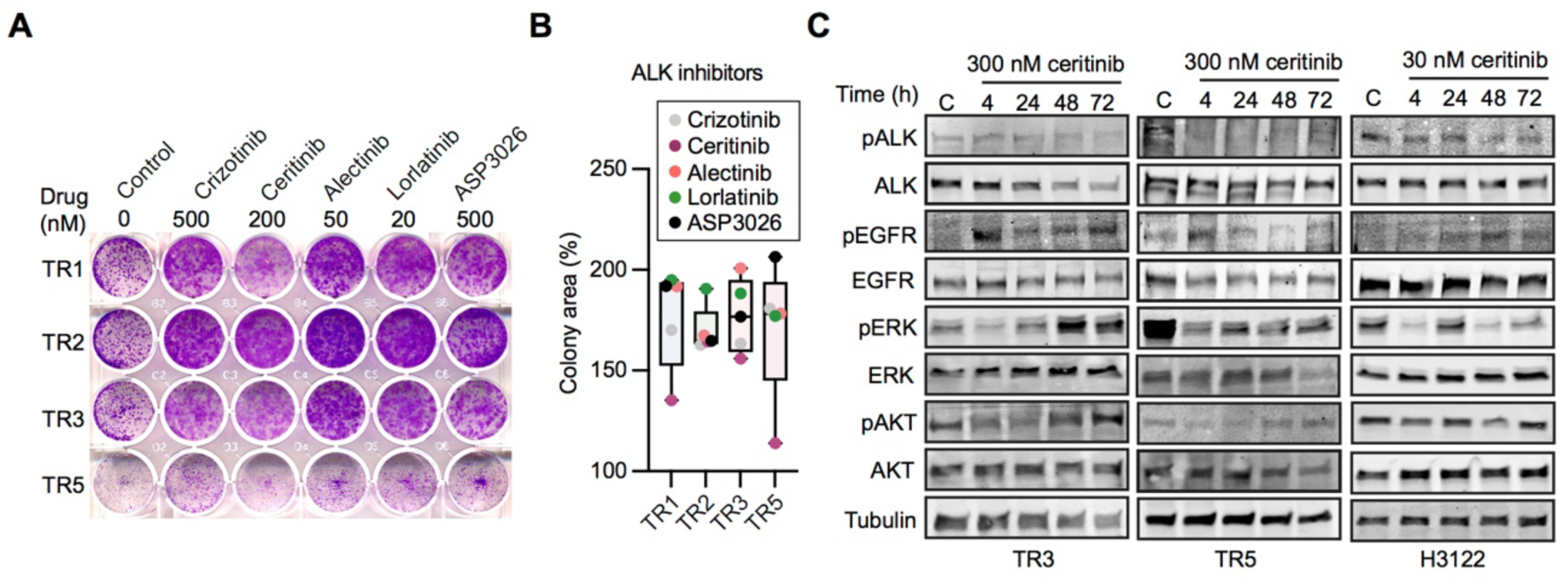
Tumor-derived cultures exhibit resistance to ALKi. (A) Representative images of clonogenicity assays of tumor-derived cultures treated for 15 days with IC50 concentrations of indicated ALKi. IC50 concentrations were derived from a three day treatment experiment. (B) Box chart showing the quantification of change in colony area with respect to vehicle control. Each dot represents an individual drug and dots are color-coded based on drug ID. (C) Immunoblots of TR3, TR5, and H3122 cells treated with vehicle (C; DMSO) and or treated with 300 nM ceritinib (TR3/TR5) or 30 nM ceritinib (H3122) for various time points (4, 24, 48, and 72 h) and probed with the indicated antibodies. ALK and phospho-ALK bands indicate variant 1 of EML4-ALK protein (117 kDa). Error bars represent minimum and maximum values.

Evaluation of MAPK and PI3K-AKT signaling in TR3, TR5, and H3122 cells treated with different ceritinib doses and times showed a substantial reduction in the level of ALK phosphorylation in all treated samples. Rebound activation of EGFR, MAPK, and PI3K-AKT pathways was consistently observed after 48 h of treatment with 300 nM ceritinib (10 nM for H3122), as evidenced by elevated phosphorylation of EGFR, ERK, and AKT (Fig. 3C). Since ALK inhibition led to adaptive activation of EGFR, and downstream signaling through MAPK and PI3K-AKT pathways, models evaluated in this study appear to exhibit EGFR-driven ALKi resistance, matching findings of other studies ^10, 17, 22^.

### A combinatorial drug screen identified PI3Kβ as a novel target to restore ALKi sensitivity

To identify treatments that may circumvent resistance to ALKi we implemented a combinatorial drug screen on tumor-derived cells using a drug panel in combination with 200 nM ceritinib. Comparing the DSSs of these drugs as a single agent or in combination with ceritinib, we identified combinatorial responses of ALKi plus EGFR inhibitors (EGFRi; n=4) and pan-ERBB inhibitors (n=12), as well as of ALKi plus PI3Kβ inhibitors (PI3Kβi; n=4) (Fig 4A). While combination responses of ALKi plus EGFRi were expected, the PI3Kβi finding was surprising, in particular because we did not detect combinatorial responses between ALKi and PI3Kα isoform inhibitors (Fig. S3A-E). To cross-validate our discoveries, we performed additional combinatorial screens in combination with gefitinib (an EGFRi) or AZD-8186 (a PI3Kβ-selective inhibitor). Both screens identified ALK as the most effective combination target (Fig 4B-C and S3E-G), altogether demonstrating that ALKi resistance in primary tumor cells was regulated by EGFR and PI3Kβ and could be effectively overcome by combined inhibition of ALK and EGFR or PI3Kβ.

**Figure 4.**
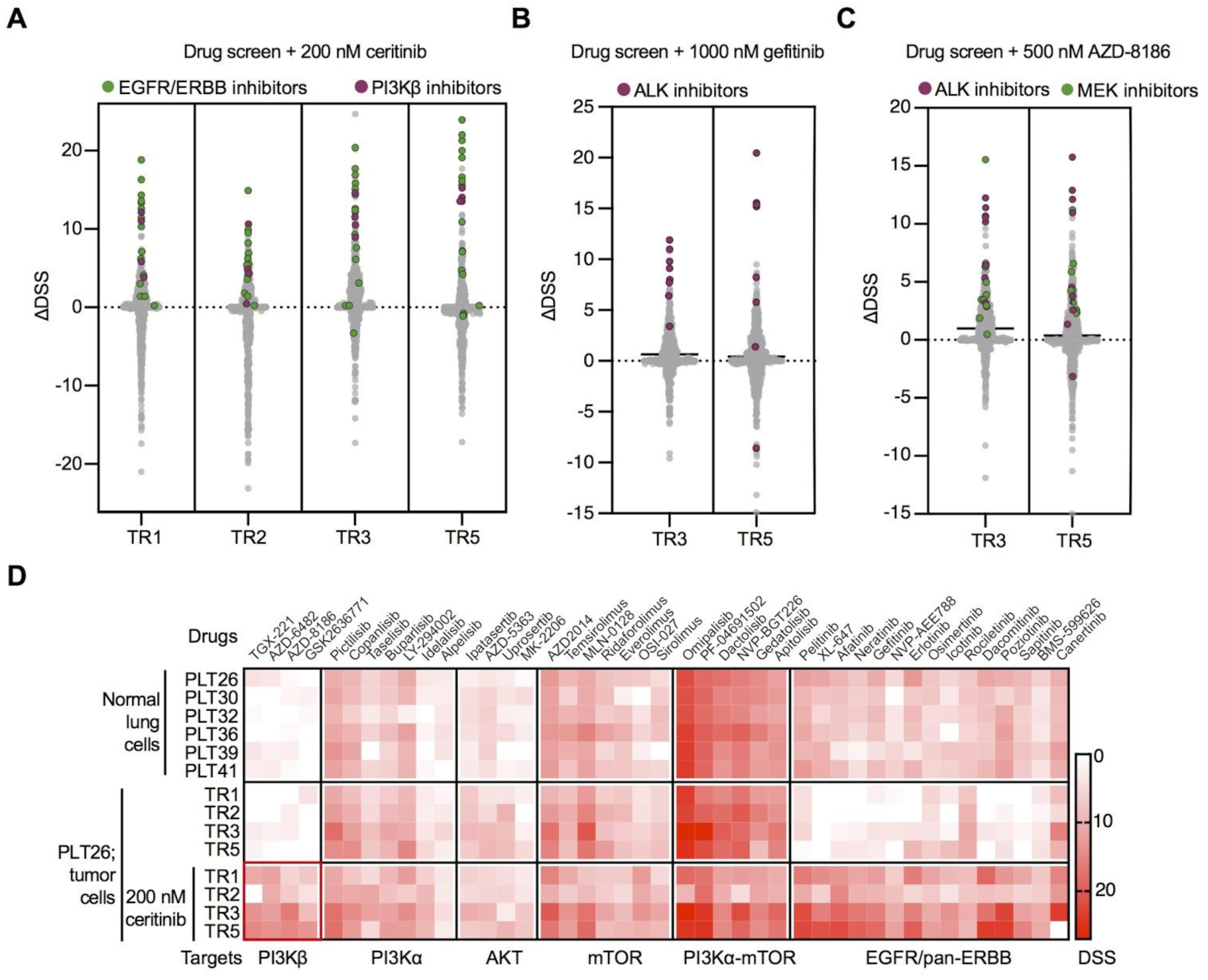
Implementation of combinatorial drug screen to identify sensitivities selective for *ALK*-rearranged lung cancer. The integrated results of a combinatorial drug screen performed on cells derived from four ALK+ tumor regions (TR1, 2, 3, and 5) or from six normal lung tissues. Note that normal lung cells PLT26 are derived from healthy tissue neighboring the ALK+ tumor. Screens were performed in combination with (A) 200 nM ceritinib (ALKi), (B) 1000 nM gefitinib (EGFRi), or (C) 500 nM AZD-8186 (PI3Kβi). (D) Heatmap showing the drug sensitivity scores of compounds inhibiting the indicated targets.

### Normal lung cells are insensitive to combined inhibition of ALK and PI3Kβ

Toxicities inflicted by anti-cancer drugs, particularly drug combinations, hinder the lab-to-clinic translation process. *Ex vivo* testing of treatments on organ-matched normal cells can provide an initial measure of generic toxicity ^23^. We utilized normal lung cells derived from six individuals to evaluate sensitivities of compounds targeting the EGFR receptor family, the PI3K-AKT-mTOR pathway, in conjunction with PI3Kα- or PI3Kβ-selective compounds. Our analysis showed that inhibition of EGFR or pan-ERBB produced generic responses in normal lung cells (Fig. 4D, S3A and S3D-E). Similarly, multiple compounds targeting various nodes of the PI3K-AKT-mTOR pathway showed toxicities in normal cells, with marked exception of compounds that selectively targeted PI3Kβ (Fig. 4D and S3B-C). Normal lung cells therefore showed no sensitivity to treatments targeting PI3Kβ, either alone or in combination with ALKi.

### Combined inhibition of ALK and PI3Kβ elicits synergistic responses in *ALK*-rearranged lung cancer cells

To widen study of the combinatorial effect of ALKi and PI3Kβi, drug response measurements were repeated and extended to two *ALK*-rearranged lung cancer cell lines, H3122 and H2228, using a wider concentration range of compounds and additionally by measuring death as a readout of drug responses. The previously detected viability-based drug sensitivities were confirmed to be synergistic in TR3 and TR5 cells, but not in H3122 and H2228 cells (Fig. 5A and S4A). However, death-based readout also amounted to a synergistic response of combinatorial ALK and PI3Kβ inhibition in H3122 cells (Fig. 5B-C and S4B). Moreover, ALKi and PI3Kβi combination treatment effectively prevented the proliferative and clonogenic potential of all *ALK*-rearranged cells (Fig. 5D-F and S4C-D).

**Figure 5.**
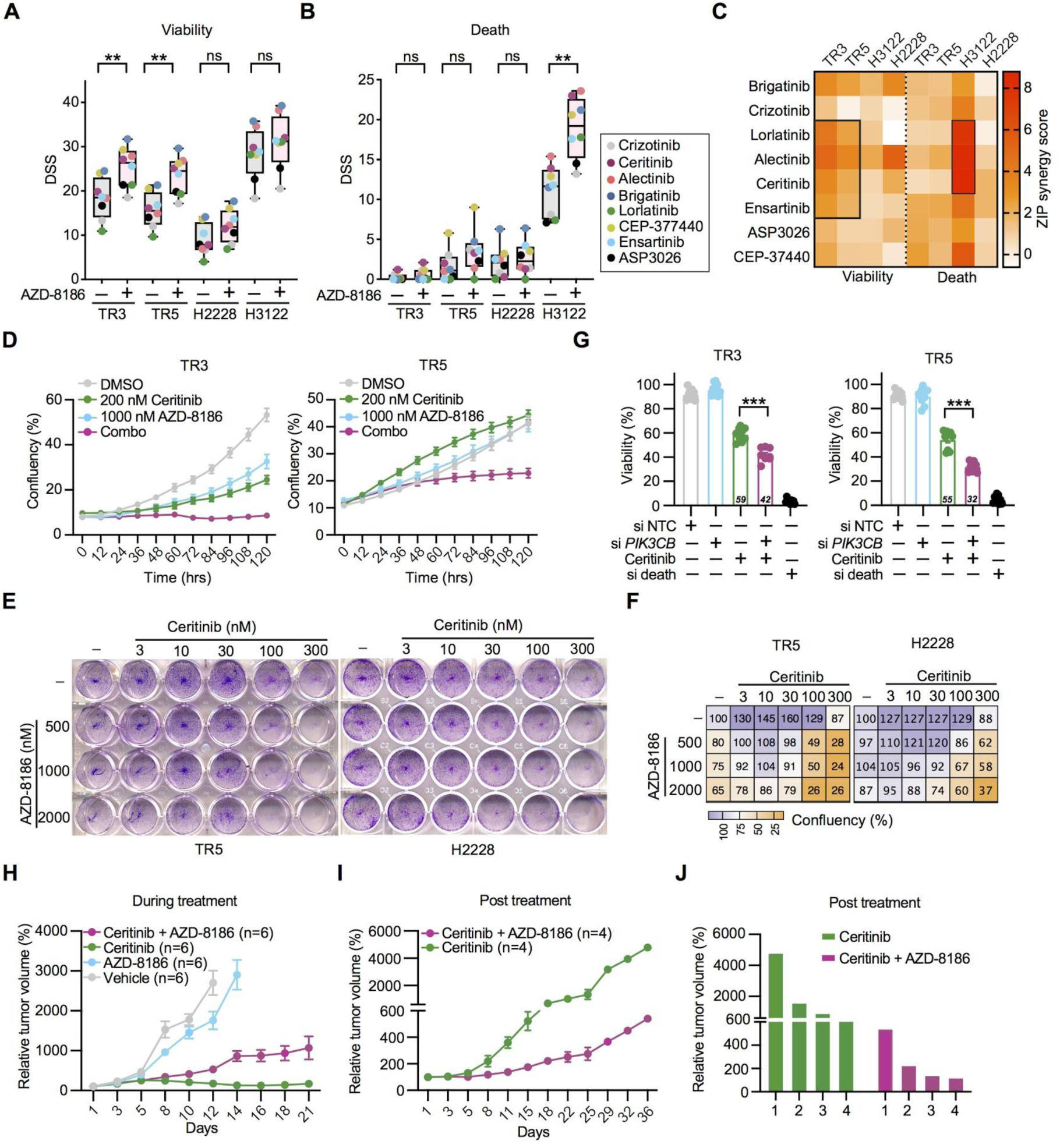
Targeting ALK in combination with PI3Kβ elicits ALK-selective responses. The DSSs of single ALKi treatment (nine doses between 0.5 and 5000 nM) or combination treatment with ALKi plus 500 nM AZD-8186. Drug sensitivities were either measured by (A) a CellTiter-Glo-based cell viability assay or (B) a CellTox Green-based cell death assay. (C) Heatmap representing zero interaction potency (ZIP)-based synergy scores calculated for each combination of ALKi plus 500 nM AZD-8186. (D) TR3 and TR5 cells were treated with vehicle (DMSO), 200 or 10 nM ceritinib, 1000 nM AZD-8186 or the combination of ceritinib plus AZD-8186; cell confluency was measured every 12 h for five days using Incucyte live cell imaging system. (E) Representative images of clonogenicity assays of TR5 and H2228 cells treated for 15 days with five different concentrations of ceritinib, three different concentrations of AZD-8186, or their combination. (F) Quantification of colony formation assays depicted in (E). (G) Percentage of viability of TR3 and TR5 cells when exposed to 200 nM ceritinib or vehicle in combination with siRNA-mediated knockdown of *PIK3CB*, non-targeting control (NTC). SiRNA and cell death siRNA were used as negative or positive controls, respectively. Viabilities of all samples were normalized to control untreated cells. Mice harboring H3122 tumors were administered with vehicle, 25 mg/kg/day ceritinib, 2×25 mg/kg/day AZD-8186, or the combination of ceritinib and AZD-8186, for 21 days. Graph shows the percentage change in tumor volume during the treatment span relative to initial tumor volume when treatment started (H), or post treatment, relative to the tumor volume when the treatment was stopped (I). (J) Waterfall plot showing percentage change in tumor volume post treatment for individual mice with the indicated treatments. Error bars represent minimum and maximum values or ± SEM. Student’s *t* test *p* values are * < 0.05, ** p <0.01, *** p <0.001.

We next performed siRNA mediated gene silencing of *PIK3CB* in TR3 and TR5 cells to verify that PI3Kβi combinatorial effects were due to selective inhibition of PI3Kβ. Despite not achieving complete knockdown of *PIK3CB* using RNAi (Fig. S4E), genetic silencing of *PIK3CB* mimicked the combinatorial responses seen with PI3Kβi (Fig. 4G). The ALKi/PI3Kβi combination did not show an effect in four *KRAS* mutant lung cancer lines, validating the selectivity of this combination for *ALK*-rearranged lung cancer (Fig. S4F). Lastly, we assessed the *in vivo* efficacy of combinatorial ALKi and PI3Kβi treatment, and treated nu/nu nude mice bearing subcutaneous tumors derived from H3122 cells with ceritinib, AZD-8186 or a combination of both treatments for 21 days. Similar to the *in vitro* findings, H3122 cell-derived tumors exhibited sensitivity to ceritinib single treatment, but while treatment discontinuation of ceritinib led to a substantial increase in tumor growth, tumors did not resume growth in the combination treatment group. This suggests that the single arm ceritinib was cytostatic, and that combinatorial ALKi and PI3Kβi treatment showed *in vivo* efficacy against H3122 ALK-rearranged cells (Fig. 5H-I). Moreover, similar to single-agent treatments, combined ceritinib and AZD-8186 treatments were well tolerated with minor or no body weight loss over the course of treatment (Fig. S4G). Altogether, these findings demonstrate that combined inhibition of ALK and PI3Kβ elicits both *ex vivo* and *in vivo* anti-tumor response and may hence offer a strategy to enhance ALKi sensitivity in *ALK*-rearranged lung cancer cells.

### Combinatorial ALKi plus PI3Kβi response appears independent of GPCR signaling and autophagy

To understand which molecular mechanisms may underpin ALKi plus PI3Kβi combinatorial effects in *ALK*-rearranged lung cancer cells, we explored the roles of autophagy and purinergic G-protein-coupled receptor (GPCR) signaling, as processes suggested to be regulated by PI3Kβ, but not PI3Kα, and previously shown to regulate ALKi resistance ^15, 17, 24, 25^. Consistent with published data ^25^, ALK inhibition triggered autophagy, and the co-targeting of ALK and PI3Kβ reduced autophagic vacuole formation (Fig S5). We therefore tested whether combining an ALKi with either autophagy inhibitors or autophagy and PI3Kα inhibitors could mimic the combinatorial cancer cell killing effect of ALKi and PI3Kβi. However, a combinatorial effect was not seen in these cases (Fig S6A-B), suggesting that PI3Kβ performs functions beyond its participation in canonical PI3K-AKT signaling and autophagy. The elevated expression of P2Y subfamily of P2 purinergic receptors was reported to be associated with ALKi resistance, both in clinical samples and cultured cells ^17^. However, inhibition of P2YRs did not enhance ALKi response in our cell systems (Fig S6C). In conclusion, our data suggest that PI3Kβ-mediated inhibition of autophagy or P2YR signaling is not sufficient to induce the combinatorial effect with ALK inhibition in *ALK*-rearranged cells.

### Resistance to ALK inhibition is associated with MAPK and PI3K-AKT pathway activation downstream of EGFR-PI3Kβ

To understand the consequences of combinatorial ALK plus PI3Kβ inhibition at the molecular level, we analyzed MAPK and PI3K-AKT pathway activities in cells treated with vehicle, ceritinib, AZD-8186, or their combination. In comparison to single treatments, cells treated with the combination showed a substantial reduction in both MAPK and PI3K-AKT activities, as well as an increase in cell death evidenced by increased PARP cleavage accompanied by reduced ERK and AKT phosphorylation (Fig 6A).

**Figure 6.**
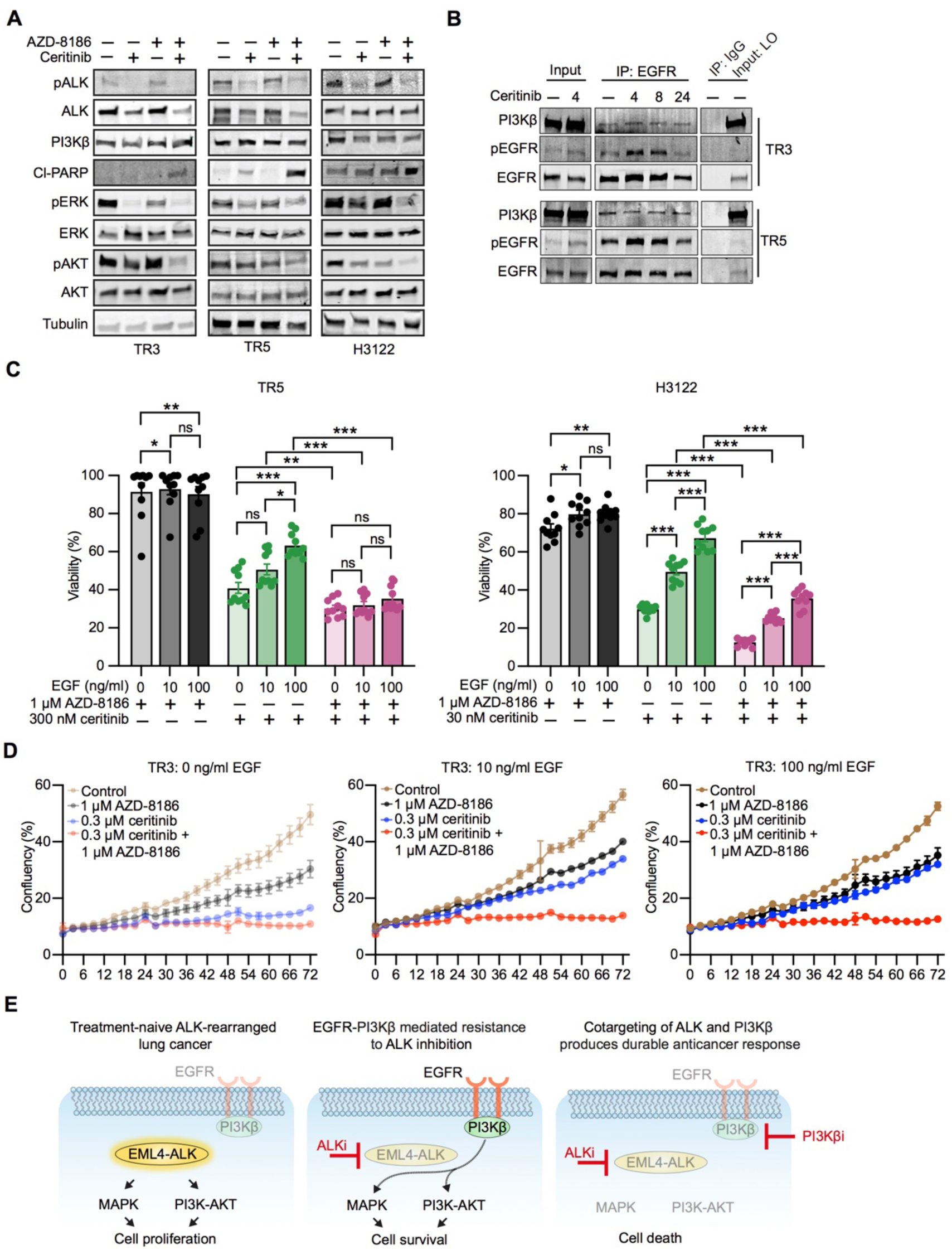
PI3Kβ contributes to cell growth and survival upon ALK inhibition in *ALK*-rearranged lung cancer. (A) Immunoblots of TR3, TR5, and H3122 cells treated with vehicle control or 300 nM ceritinib (TR3/TR5) or 30 nM ceritinib (H3122) or 1000 nM AZD-8186 or their combination for 72 h and probed with the indicated antibodies. (B) TR3 and TR5 cells were treated with 300 nM ceritinib for different time points (4, 8, and 24 h) and lysates were immunoprecipitated with EGFR antibody. Precipitates were analyzed by immunoblotting and probed with the indicated antibodies. (C) TR5 and H3122 cells were co-treated with different doses of EGF (0, 10, and 100 ng/ml) and ceritinib or AZD-8186 or their combination for 72 h. Percentage viabilities of drug-treated cells were normalized to cells co-treated with different doses of EGF (0, 10, and 100 ng/ml) or DMSO. (D) TR3 cells were co-treated with different doses of EGF (0, 10, and 100 ng/ml), vehicle (DMSO), ceritinib, AZD-8186, or the combination of ceritinib plus AZD-8186; cell confluency was measured every 3 h for 72 h using the Incucyte live cell imaging system. (E) Graphical illustration describing resistance to ALKi in *ALK*-rearranged lung cancers in which tumor cells are dependent on EGFR or PI3Kβ for cell growth and survival.

Next, we explored how PI3Kβ function is regulated upon ALK inhibition. Considering that the ERBB family member ERBB3 was reported to recruit PI3Kβ and thereby drive PI3Kα inhibitor resistance in *HER2*-amplified and *PIK3CA* mutant cancers ^18^, we hypothesized that ALK inhibition may similarly activate EGFR and thereby regulate PI3Kβ function. We therefore immunoprecipitated EGFR from TR3 and TR5 cell lysates treated with DMSO control or 300 nM ceritinib for different times and assessed the phosphorylation of EGFR and PI3Kβ binding. EGFR phosphorylation was confirmed to significantly increase following ALK inhibition (Fig 6B). Interestingly, PI3Kβ co-precipitated with EGFR, demonstrating that EGFR physically interacts with PI3Kβ in those cells (Fig 6B). Moreover, treatment of TR3, TR5, H3122, or H2228 cells with EGF elicited increased resistance to ALK inhibition, while the combined inhibition of ALK and PI3Kβ rescued the EGF-induced resistance, together suggesting that inhibition of PI3Kβ restricts EGFR-mediated bypass resistance to ALK inhibition in *ALK*-rearranged lung cancer (Fig 6C-D and S7A-B).

In summary, dissection of the mechanisms underpinning combinatorial ALK and PI3Kβ inhibition efficacy suggested that the sensitization of *ALK*-rearranged cancer cells to ALK inhibition by selective PI3Kβ inhibition was, at least in part, mediated through a blockade of EGFR-mediated rebound activation of MAPK and PI3K-AKT signaling, enhancing cell death (Fig 6E).

## DISCUSSION

Clinical response to ALKi tends to be short-lived due to the acquisition of ALK-independent drug resistance ^2^. We here studied primary cells derived from an aggressive *TP53* mutant and chemoresistant *ALK*-rearranged tumor that exhibited region-specific EMT features to identify novel/effective combination treatments to treat *ALK*-rearranged NSCLC. Tumor cultures derived from the region that exhibited a mesenchymal phenotype grew as cell clusters and expressed vimentin, and, consistent with earlier findings ^26, 27^, were less responsive to treatment with ALKi than cells from epithelial phenotype regions of the tumor. We further identified selective PI3Kβ inhibition as a promising novel therapeutic target to potentiate ALK inhibition responses in *ALK*-rearranged lung cancer. We demonstrate that the co-targeting of ALK and PI3Kβ leads to blockade of both MAPK and PI3K-AKT signaling, attenuates EGFR mediated adaptive resistance, and elicits tumor cell-selective toxicity with efficacy even in *TP53* mutant *ALK*-rearranged cells that have undergone EMT. Our study therefore demonstrates the utility of primary patient tumor-derived cultures for pharmacogenomic profiling and dissection of drug resistance mechanisms.

Both the α and β PI3K class IA isoforms are ubiquitously expressed in both normal and malignant tissues, but these subunits are regulated differently, induce different modes of signaling, and have divergent roles in health and disease ^28^. The PI3Kα isoform is a well-established oncogenic driver that acts downstream of RTK signaling that is frequently oncogenically mutated and constitutes a drug target for several cancer types, including lung cancer ^28^. On the other hand, PI3Kβ has both kinase-dependent and -independent functions as an effector of both RTKs and GPCRs ^15^. Similar to PI3Kα, PI3Kβ participates in canonical PI3K-AKT signaling, but also uniquely regulates DNA replication, cell cycle progression, nuclear envelope maintenance and nuclear pore complex assembly, autophagy, and apoptosis^15^. Despite PI3Kβ rarely being mutated in cancers, PI3Kβ expression correlates with poor prognosis in early-stage NSCLC patients ^29^. It has also been proposed as a drug target in PTEN-deficient cancers ^30^. In this respect our discovery of PI3Kβ as a target to enhance ALKi sensitivity in *ALK*-rearranged lung cancer is unexpected, as combinatorial PI3Kβi sensitivity is independent of *PTEN/PIK3CA* mutations or resistance to PI3Kα inhibitors. Since PI3Kβ is a downstream node for signaling directed by those RTKs that are widely implicated in driving resistance to ALKi, particularly IGF1R ^19, 20^, ERBB2/3 ^17, 18^ and GPCRs ^16, 17^, targeting PI3Kβ may provide a common target to counter a variety of bypass/acquired ALKi resistance mechanisms. Hence, our findings warrant further testing of the efficacy of PI3Kβi drug combinations on a larger set of patient-derived samples.

Previous research has identified various bypass signaling tracks that can mediate ALKi resistance, fostering the formulation of combinatorial treatment options that may overcome or even prevent drug resistance. The efficacy and tolerability of co-targeting ALK with MEK (NCT03202940, NCT03087448) or mTORC1 (NCT02321501) are currently tested in patients with *ALK*-rearranged NSCLC. In addition, following promising preclinical findings, the clinical testing of combinations co-targeting ALK with AXL, SRC, KIT, MET or SHP2 are anticipated in the near future. Eventually, only those combinations that exhibit both efficacy and manageable toxicity profiles will be approved. It is therefore notable that the co-inhibition of MEK, mTOR, SRC, KIT, MET or SHP2 did not enhance the ALKi response in our study. Use of organ-matched healthy epithelial cells may help to exclude treatments that show generic toxicity ^23^. We here for example detected sensitivity to HSP90 inhibition in both normal epithelial and cancer cell cultures, corroborating the finding that HSP90 inhibition induces toxicity without tumor-selective response in patients carrying *ALK*-rearranged NSCLC ^31, 32^. Likewise, and in spite of the fact that compelling preclinical data support the co-targeting of ALK and EGFR, we found that normal lung cells showed generic toxic response to co-targeting EGFR, which aligns with detected severe toxicity in NSCLC patients receiving combinatorial treatments with EGFRi ^33, 34^. Based on the evidence presented here, it appears that EGFR and PI3Kβ might share common pathways towards resistance, indicating that co-targeting of ALK plus PI3Kβ may be broadly applicable as a strategy to counteract EGFR-mediated adaptive resistance to ALKi also in other tumors and in other cancer types. Promisingly, PI3Kβ-selective inhibitors are being investigated in seven phase-I/II clinical trials as a single agent or in combination with chemo- or immunotherapy (Table S3), following initial reports of satisfactory tolerability and efficacy ^35, 36^.

From a heterogeneity standpoint, varied expression of lung adenocarcinoma markers, EMT, and tumor vasculature was observed across six different regions of the same tumor, suggesting that a single tumor biopsy would not provide a holistic assessment of this particular *ALK*-rearranged cancer. Validating our approach, cells derived from the tissue region that predominantly exhibited a mesenchymal phenotype showed the lowest sensitivity to ALK inhibition, matching published data ^26, 27^. Given that cells derived from all tumor regions carried the same *TP53* mutation, representing both epithelial and mesenchymal phenotypes, as well as established *ALK*-rearranged H3122 (*TP53* mutant) and H2228 (*TP53* and *NFE2L2* mutant) cells showed sensitivity to the ALKi plus PI3Kβi combination, our findings suggest that this combination carries promise to counteract ALKi resistance associated with EMT, *TP53* or *NFE2L2* mutations ^2, 21^.

To understand how co-targeting PI3Kβ improved ALK inhibitory responses we explored involvement of EGFR and GPCR signaling as well as autophagy as these processes are linked to ALKi resistance and PI3Kβ functions ^16, 17, 24, 25^. While the activation of purinergic GPCRs was previously shown to drive ALKi resistance ^17^, purinergic GPCR inhibition did not improve the ALKi response in cells cultured from our clinical sample. Hence, given that inhibition or activation of EGFR respectively increased or decreased ALKi responses, it appeared that EGFR rather than GPCRs regulated PI3Kβ in the cells studied here. Interestingly, we measured a previously unreported interaction between EGFR and PI3Kβ, fitting with the finding that blockade of EGFR-PI3Kβ signaling synergizes with ALK inhibition. Moreover, we observed activation of autophagy as a result of ALK inhibition, which aligns with previous studies ^25^, and we extend this data by demonstrating attenuation of ALKi-induced autophagy by PI3Kβ inhibition. Nevertheless, autophagy inhibitors did not mimic PI3Kβ inhibition in eliciting combinatorial ALKi response, suggesting that PI3Kβ exerts functions beyond autophagy in conferring survival following ALK inhibition.

In summary, our study proposes PI3Kβ as a novel regulator of ALKi resistance in *ALK*-rearranged lung cancer cells, showing efficacy even in aggressive *TP53* mutant and mesenchymal cells. Given that multiple salvage signals are activated upon resistance to ALKi, the co-targeting of PI3Kβ may provide a promising therapy option as PI3Kβ acts as a common effector of multiple salvage pathways, including EGFR, autophagy and GPCRs. Importantly, we found that co-targeting of PI3Kβ showed reduced toxicity in normal lung epithelial cells when compared with co-targeting of EGFR, possibly providing for a relatively safe clinical direction to treat *ALK*-rearranged tumors.

## MATERIAL AND METHODS

### Culture of patient-derived cells and cell lines

Procedures conducted in this study were performed in accordance with licenses approved by the Coordinating Ethics Committee of the University of Helsinki (85/13/03/00/2015, HUS-1204-2019). Clinical samples were collected at the Helsinki University Central Hospital (HUCH) with the patient’s written informed consent. Immediately after surgery, a thoracic pathologist dissected the tumor and tumor-adjacent normal lung tissues. Tissue pieces were manually minced using a sterile scalpel, and then enzymatically digested using the Tumor Dissociation Kit (Miltenyi, 130-095-929) and gentleMACS Dissociator (130-093-235), following the manufacturer’s instructions. Single cell suspensions were further processed using a Miltenyi’s human EpCAM (CD326) cell isolation kit (130-061-101) to enrich EpCAM+ cells. Both normal and tumor EpCAM+ cells were cultured in the presence of irradiated (30 Gy) 3T3 feeder cells in F-medium (see Supplementary Method for composition) using Conditional Reprogramming (CR) culture protocol as described in ^23^. Feeder 3T3 cells were cultured in DMEM (Thermo Fisher Scientific, 42430-025) supplemented with 10% FBS (Gibco, 10270-106). Differential trypsinization was utilized for culture propagation when cells reached 80% confluency ^37^. Lung cancer cell lines (H3122, H2228, A549, H460, Calu-1, and H1437) were cultured in RPMI-1640 (Lonza, 15-1675) supplemented with 10% FBS. All cultures were maintained in a humidified incubator at 37 °C with 5% CO2. H3122 (Cellosaurus accession ID: CVCL_5160), H2228 (CVCL_1543), A549 (CVCL_0023), H460 (CVCL_0459), Calu-1 (CVCL_0608), and H1437 (CVCL_1472) cells were purchased from the ATCC and STR-profiled at the Genotyping Unit of Technology Centre at the Institute for Molecular Medicine Finland (FIMM).

### Drug Sensitivity and Resistance Testing (DSRT) assay

DSRT assays for anticancer compounds as a single agent or in combination with other compounds were performed as previously described ^38^. For drug screening, 384-well plates (Corning; 3712) with compounds were prepared in advance by dispensing compounds using an Echo 550 liquid handler (Labcyte), at five concentrations covering a 10,000-fold concentration range. For storage, the pre-drugged plates were kept in pressurized StoragePods (Roylan Developments Ltd.) under inert nitrogen gas. For drug screening, the pre-drugged compounds were dissolved in 5 µl of culture medium per well, with (1:2,000 final volume) or without CellTox Green (Promega) depending on the experiment, and 20 µl cell suspension per well was seeded at a concentration of 1500 cells/well. After 72 h incubation at 37 °C, cell death was assessed by measuring fluorescence signals from CellTox Green (485/520 nm excitation/emission filters). Subsequently, to assess cell viability, 25 µl/well CellTiter-Glo reagent (Promega) was added and luminescence was recorded. Both fluorescence and luminescence readouts were recorded using a PHERAStar FS plate reader (BMG Labtech). To plot dose-response curves for each drug, the Marquardt-Levenberg algorithm was implemented using the in-house bioinformatic ‘Breeze’ pipeline ^39^. To compare drug responses across samples, Drug Sensitivity Scores (DSSs) were calculated using dose–response curve parameters including the IC50, slope, top, and lower asymptotes, as described ^40^. To manually assess the drug responses of single agents or drug combinations, a denser concentration range (nine doses between 0.5/1 and 5,000/10,000 nM) of compounds was used. Cells (1,500 per well) were seeded in 384-well plates in 20 µl of media. After 24 h incubation at 37°C, cells were treated with vehicle control or drug in 10 µl of media with three technical replicates for each condition. After 72 h incubation at 37°C, cell viability was quantified using CellTiter-Glo reagents. The relative cell viability was calculated using the formula: (cell viability of drug treatment) / (cell viability of vehicle control)×100.

### Immunoblotting

Lysates were prepared from tumor tissue or cultured cells using RIPA buffer supplemented with fresh protease and phosphatase inhibitors (Roche). Protein quantification was performed using the BCA Protein Assay (G Biosciences; 786-570). Lysates were fractionated using Mini-PROTEAN TGX precast gels (Biorad) and transferred to PVDF membranes (Millipore, IPFL00010) using the XCell II blot module (Thermo-Scientific). After transfer, membranes were blocked for 30 min at room temperature using Odyssey Blocking Buffer. Two-color immunoblotting was performed using Odyssey Blocking Buffer (LI-COR) and IRDye (800CW/680RD) secondary antibodies (LI-COR) diluted 1:10,000 in Odyssey Blocking Buffer. Membranes were scanned using an Odyssey infrared imager (LI-COR) and quantifications were performed using Image Studio software (LI-COR). Primary antibodies are listed in the Supplemental Experimental Procedures.

### Co-immunoprecipitation

Co-immunoprecipitation (Co-IP) experiments were performed using the Thermo Fisher Scientific Pierce Co-IP Kit (26149), following the manufacturer’s protocol. In brief, cells were exposed to drugs when cell confluency reached 60-70%. After 4 or 24 h of drug exposure, cells were lysed with an ice-cold Lysis/Wash Buffer supplemented with fresh protease and phosphatase inhibitors (Roche). Cell lysates were pre-cleared by incubating the lysate with Control Agarose Resin at 4°C for 1 h. To prepare antibody-immobilized AminoLink Plus Coupling Resin, 25 µl Coupling Resin, and 1 µg of rabbit anti-EGFR (Santa Cruz Biotechnology; sc-120) or IgG control antibody (Invitrogen; 02-6102) were co-incubated in a spin column for 2 h at room temperature. Columns were then washed twice with 1X Coupling Buffer and six times with Wash Solution, and centrifuged after each wash. Subsequently, pre-cleared cell lysate was added to the column containing antibody-coupled resin, and incubated overnight at 4°C with gentle mixing. Proteins were eluted using 60 µl of Elution Buffer, and eluates were analyzed by western blotting with primary antibodies for EGFR, pEGFR, PI3Kβ, and tubulin. See Supplementary Information section for antibody details.

### IncuCyte confluency assay

Cells were seeded at a density of 5,000–10,000 cells per well in 96-well plates and were exposed to drugs or DMSO on the following day. Immediately after drug treatment, cells were followed by live-cell imaging using an IncuCyte ZOOM microscope (Essen Bioscience) by taking pictures every 3 or 4 h for a total of 72 or 120 h. Images were then quantified for cell confluency (cell surface area coverage as confluence values) using the IncuCyte application software, and cell confluency data were plotted with GraphPad Prism.

### Colony formation assay

Cells were seeded at a density of 10,000–20,000 cells per well in 24-well plates and were exposed to drugs or DMSO on the following day. Medium and DMSO/drugs were replaced every 72 h for 15 days. Cells were fixed with colony fixation solution (acetic acid/methanol 1:7 (vol/vol)) and stained with 0.5% crystal violet. After drying, stained plates were scanned with the Cytation 5 Cell Imaging Multi-Mode Reader (BioTek) and the number and size of colonies were quantified with the compatible Gen5™ Multi-Mode Reader and Imager Software.

### Autophagy analysis

Cells were seeded at a density of 10,000–20,000 cells per well in 96-well plates and exposed to drugs or DMSO on the following day. Following 24 h incubation, cells were co-stained with CYTO-ID Green detection reagent and Hoechst 33342 nuclear stain according to the manufacturer’s instructions (Enzo Life Science). Cells were observed and imaged using Opera Phenix High-Content Screening System (PerkinElmer). Quantification of the number of autophagic vacuoles, vacuoles per cell, their size, and the intensity of the vacuoles was carried out using the system’s Harmony software.

### *In vivo* treatment experiment

Treatment studies on H3122 xenografts were performed at the Charles River facilities. This study was carried out in strict accordance with the recommendations in the Guide for the Care and Use of Laboratory Animals of the Society of Laboratory Animals (GV SOLAS) in an AAALAC accredited animal facility. All animal experiments were approved by the Committee on the Ethics of Animal Experiments of the regional council (Regierungspräsidium Freiburg, Abt. Landwirtschaft, Ländlicher Raum, Veterinär-und Lebensmittelwesen - Ref. 35, permit-#: G-18/12). Both male and female NSG mice of 6-8 old weeks were injected subcutaneously with 2 × 10^6^ cells in 50% Matrigel. Once average tumor volume reached 100 mm^3^, animals were randomized to different treatment groups (n = 6 per treatment group) and were administered with vehicle control, ceritinib alone, AZD-8186 alone or combination of ceritinib plus AZD-8186. Ceritinib was administered at 25 mg/kg body weight p.o. daily for 21 days. AZD-8186 was administered at 2 × 25 mg/kg body weight p.o. daily for 21 days. Ceritinib (25 mg/kg) and AZD-8186 (2 × 25 mg/kg) were administered together p.o. daily for 21 days. The control vehicle (0.5% Methylcellulose, 0.5% Tween80) p.o. for 21 days. Tumors were measured using electronic calipers twice a week, and tumor volumes were calculated using the formula length × width^2^ × 0.52. Body weights were recorded in parallel to the tumor volume measurement. Animals were daily monitored for signs of morbidity and/or mortality.

### Drug treatment with or without EGF

Cells (5,000 cells/well) were cultured in 96-well plates in media without EGF. After 24 h, cells were treated with indicated drug concentration or vehicle control in media supplemented with 10 or 100 ng/ml fresh EGF, or without addition of EGF. Drug and EGF treatments were repeated every 24 h for 3 days. After 72 h of total drug treatment, cell viability was quantified using CellTiter-Glo reagents. The relative cell viability was calculated using the formula: (cell viability of drug treatment) / (cell viability of vehicle control)×100.

### Statistics and reproducibility

GraphPad Prism 9 (GraphPad Software Inc) was used to generate all presented figures presented and to perform statistical analyses of experimental data. Statistical significance was assessed using a Student’s t test and nonparametric Mann-Whitney test. P-values greater than 0.05 were considered as statistically significant. Error bars indicate standard deviation or standard error of the mean. Pearson’s correlation coefficients were used to assess the significance of correlations and displayed in the XY plots. DSRT, NGS, and IHC experiments were performed a single time, with biological replicates (n=2-6). Immunoblot analyses including co-immunoprecipitation, FISH, NRF2-related signatures, and validation of siRNA mediated *PIK3CB* knockdown were performed two times, with similar results. Follow-up experiments including colony formation assay, IncuCyte confluency assay, CYTO-ID based autophagy assay, and experiments evaluating the effect of *PIK3CB* knockdown, EGF treatment, GPCR inhibition or autophagy inhibition on responses of ALKi, PI3Kβi or combination of ALKi plus PI3Kβi were performed three times, with similar results. Xenograft experiments were performed once at Charles River by different experimenters.

## Data availability

The authors declare that the main data supporting the findings of this study are available within the article and its Source Data File. The data presented in Fig S2F of this study are available from the GDSC database portal (https://www.cancerrxgene.org/). Other data, such as IHC images midst sharable following GDPR regulations is available from the corresponding author upon request.

## Supplementary Information

Table of contents:

## Supplementary Methods

- F-medium
- Immunohistochemistry
- Fluorescence In Situ Hybridization (FISH)
- Genetic analysis
- RNA extraction, cDNA synthesis, and qRT-PCR
- *PIK3CB* RNAi and ALKi drug screens
- Table S1. Details of primary antibodies used in immunohistochemistry and western blotting analyses
- Table S2. Drug library used for Drug Sensitivity and Resistance Testing
- Table S3. Clinical trial information for PI3Kβi

## Supplementary Figure Legends

- **Figure S1.** Tumor-derived cultures retain genotypes and phenotypic features of tumors
- **Figure S2.** Ceritinib treatment leads to reactivation of ERK, AKT, and EGFR
- **Figure S3.** Normal lung cells are insensitive to combined inhibition of ALK and PI3Kβ
- **Figure S4.** The PI3Kβi AZD-8186 increases ceritinib efficacy in *ALK*-rearranged lung cancer
- **Figure S5.** ALK inhibition activates autophagy
- **Figure S6.** Inhibition of autophagy or GPCR does not improve the response of ceritinib
- **Figure S7.** Combined inhibition of ALK and PI3Kβ overcomes EGFR mediated resistance in *ALK*-rearranged lung cancer cells

## F-medium

F-medium is a combination of 1:3 v/v DMEM : F-12 nutrient HAM supplemented with 5% FBS, 10 ng/ml EGF (BD Biosciences; 354052), 5 µg/ml insulin (Sigma; I2643), 24 µg/ml adenine (Sigma; A2786), 0.4 µg/ml hydrocortisone (Sigma; H4001), 10 ng/ml cholera toxin (List Biological laboratory; 100B), and 10 µM ROCK inhibitor (Y-27632; ENZO) and penicillin-streptomycin (Gibco; 15140-122) to final concentrations of 100 Units/ml and 100 µg/ml, respectively.

## Immunohistochemistry

Tissue processing and IHC procedures were performed essentially as described previously ^41^. Antibodies and stain-specific details are listed in the Supplemental Information section. To acquire whole slide scans of stained tissue sections the Pannoramic 250 digital slide scanner (3DHISTECH) was used and the scanned TIFF images are exported using the Pannoramic Viewer (3DHISTECH).

## Fluorescence In Situ Hybridization (FISH)

The *ALK*-rearrangement status of the samples was evaluated by FISH using *ALK* dual-color break-apart probe according to the manufacturer’s protocol (Vysis, Abbott Molecular Inc.). Tumor cells nuclei with split red and green signals were defined as positive for *ALK-* rearrangement, and at least 100 cells were evaluated for each sample to conclude the *ALK*-rearrangement status. Samples displaying *ALK*-rearrangement in more than 10% of the cells were labeled as positive.

## Genetic analysis

Genomic DNA was extracted from healthy lung and tumor tissue samples and from corresponding CR cultures using a DNeasy Blood & Tissue kit (Qiagen). Targeted next-generation sequencing was performed using the NimbleGen Cancer Panel (captures the exons of 578 cancer-related genes) and the Illumina HiSeq2500 system in HiSeq high output mode using v4 chemistry or HiSeq Rapid run mode using v2 chemistry (Illumina), as described ^23^. Variants were removed if the variant allele frequency was < 2%; tools employed for variant calling have been outlined earlier ^42^.

## RNA extraction, cDNA synthesis, and qRT-PCR

Total RNA was extracted using NucleoZOL (Macherey-Nagel) following the manufacturer’s instructions. One microgram of the RNA was reverse transcribed to cDNA using the Transcriptor First Strand cDNA Synthesis Kit (Roche) according to the manufacturer’s protocol. cDNA templates were assayed in 10 µl PCR reactions with 10 nM of each primer and 10 ng of template cDNA with LightCycler 480 Real-Time PCR system (Roche) according to the protocol of Fast Start Universal Probe Master (Roche). mRNA levels were determined using Universal Probe Library System (Roche). Specific primer and probe information is listed in Table S3. For analysis, Roche LC Software was used for Cp determination (by the second-derivative method). β-actin was used for normalization.

## *PIK3CB* RNAi and ALKi drug screens

*PIK3CB* gene silencing was achieved by RNA interference using a reverse transfection procedure. 384-well plates (Corning; 3712) with siRNAs were prepared in advance by dispensing siRNAs using an Echo 550 liquid handler (Labcyte). Plates were either used immediately or foil-sealed and stored at −20°C until use. ON-TARGET plus SMART pool siRNA (Dharmacon) against *PIK3CB* was used (10, 20 or 40 nM). Before transfection, siRNA molecules were reconstituted in OptiMEM (5 µl/well) medium (Invitrogen) containing Lipofectamine RNAiMAX (25 nl/well; Life Technologies) using a Multidrop™ Combi nL Reagent Dispenser (Thermo Fisher Scientific). After 30 min incubation at RT on an orbital shaker, 20 µl cell suspension per well was seeded in the siRNA plates. After 24 h incubation at 37°C, cells were treated with DMSO or 200 nM ceritinib and further incubated for 72 h at 37°C. Cell viability was measured using CellTiter-Glo reagent as per the manufacturer’s instructions. Cells transfected with 10 nM of non-targeting siRNA (Qiagen) or AllStars Hs Cell Death Control siRNA (Qiagen) served as a negative and positive control. To verify the knockdown of *PIK3CB*, 1×10^5^ cells were seeded in each well of a six-well plate, after 24 h, cells were transfected with 10 nM *PIK3CB* siRNA using Lipofectamine RNAiMAX according to the manufacturers’ instruction. At 72 h after transfection, lysates were collected and subjected to immunoblotting analyses.

## Author Contributions

SST, KW and EWV conceived and designed the study; SST performed *in vitro* experiments, analyzed the data, and generated the figures; MIM performed clinical pathology analyses; JS designed and performed the *in vivo* xenograft experiment; NL and AH implemented IHC staining and scanned slides; WS advised on the study design; SA characterized cell lines for NRF2-related signatures; JR performed surgeries, and JR, AK, and MIM collected clinical data, received patients’ informed consent, and managed the primary tissue workflow; SST, KW, and EWV wrote the manuscript; MIM, JS, WS, and ALL gave critical comments and corrections to the manuscript; EWV, JS, ALL and KW supervised the study.

## Acknowledgements

We dedicate this work to the patients who supported our research by consenting access to clinical specimens. We are thankful to Merja Räsänen for processing patient consents, Astrid Murumägi for reagents and guidance on the establishment of the CR culture protocol and Tarja Salonen from HUSLAB Genetics Laboratory (HUS Diagnostic Center) for ALK-rearrangement FISH analysis. We thank Soili Kytölä, the Sequencing Core Facility at FIMM, HiLIFE, at the University of Helsinki and the Laboratory of Genetics, HUS Diagnostic Center, HUSLAB, at the Helsinki University Hospital for NGS analysis. We thank the thoracic pathologists who helped with the clinical specimen collection, the FIMM Digital Microscopy and Molecular Pathology Unit for scanning histological slides, the FIMM High-Throughput Biomedicine facility for drug screening resources, and the FIMM High Content Imaging and Analysis unit for imaging and data analysis for autophagy evaluation. We thank Antti Hassinen for providing assistance for imaging and data analysis. We thank members of the Verschuren and Wennerberg labs for support and guidance. The study received financial support from the University of Helsinki Integrative Life Science doctoral program (SST); Biomedicum Helsinki Foundation (SST); Cancer Foundation Finland (SST), K. Albin Johansson Foundation (SST), Genetics and Mechanisms in Translational Medicine (GenomMed) Doctoral programme (SA, Marie Skłodowska Curie grant agreement No 740264), HUSLAB and the Finnish Medical Foundation (MIM); the Academy of Finland (EWV; grant 307111, ALL grant 332697); Novo Nordisk Foundation (KW; Novo Nordisk Foundation Center for Stem Cell Biology, DanStem; grant no NNF17CC0027852); the Sigrid Jusélius Foundation (KW, ALL); and a HiLIFE/University of Helsinki HiPOC 2020 grant (EWV).

---

**Figure S1.**
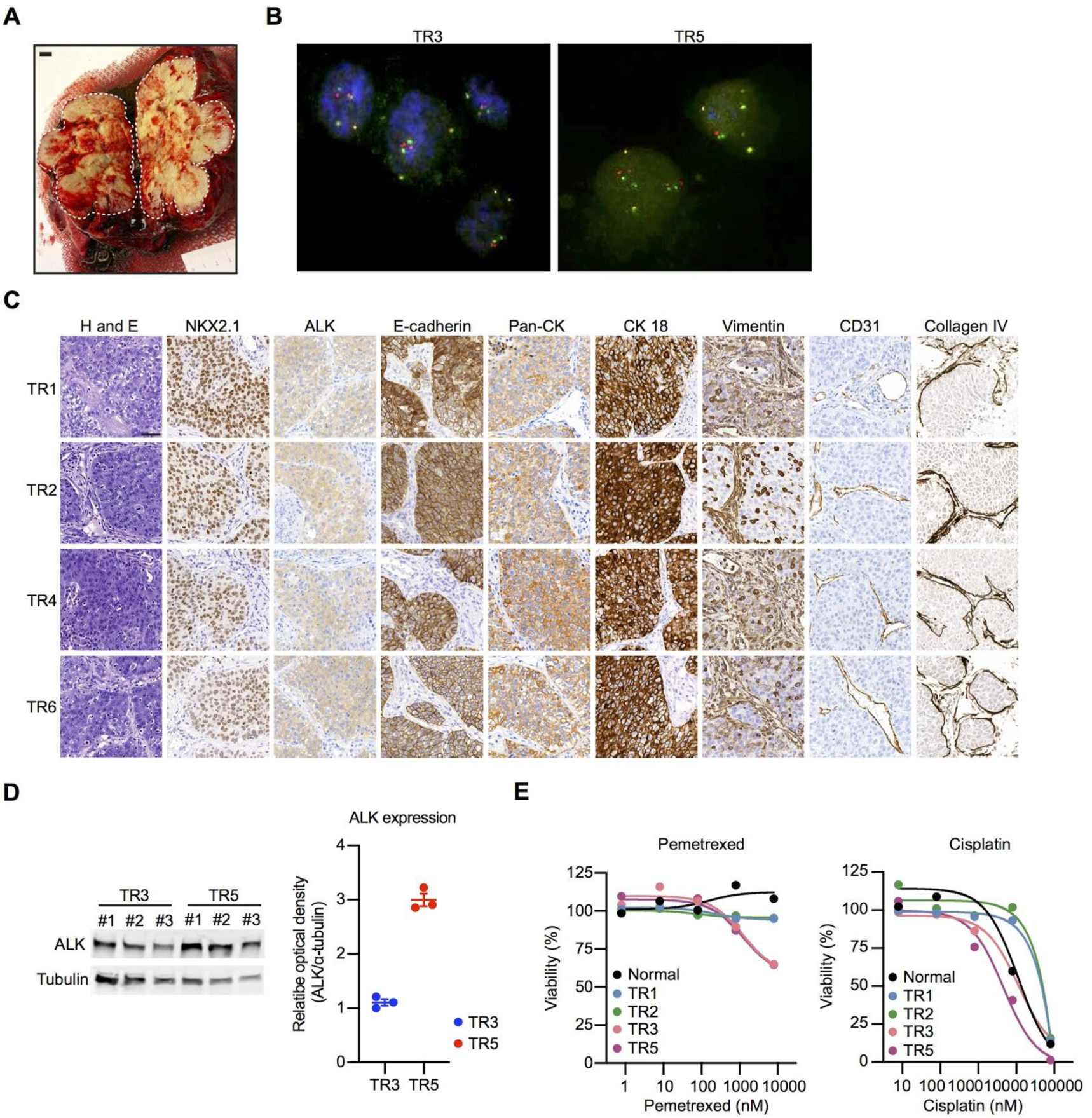
Tumor-derived cultures retain genotypes and phenotypic features of tumors. (A) Image of the surgically removed tumor tissue used to collect samples from multiple regions. The scale bar corresponds to one cm. (B) FISH analysis of TR3- and TR5-derived cells for ALK rearrangement. (C) Extended dataset for Figure 1B. Representative images of hematoxylin and eosin (H&E) and IHC staining performed on different regions of the *ALK*-rearranged tumor tissue. The scale bar corresponds to 50 µm. (D) Immunoblots (left) of *ALK*-rearranged tumor-derived cultures and probed with the indicated antibodies. Quantification (right) of relative expression of ALK in TR3 and TR5 cells. (E) Dose-response curves of normal lung or tumor-derived cells treated with pemetrexed and cisplatin; these were chemotherapy drugs given to the patient. Error bars represent ± SEM.

**Figure S2.**
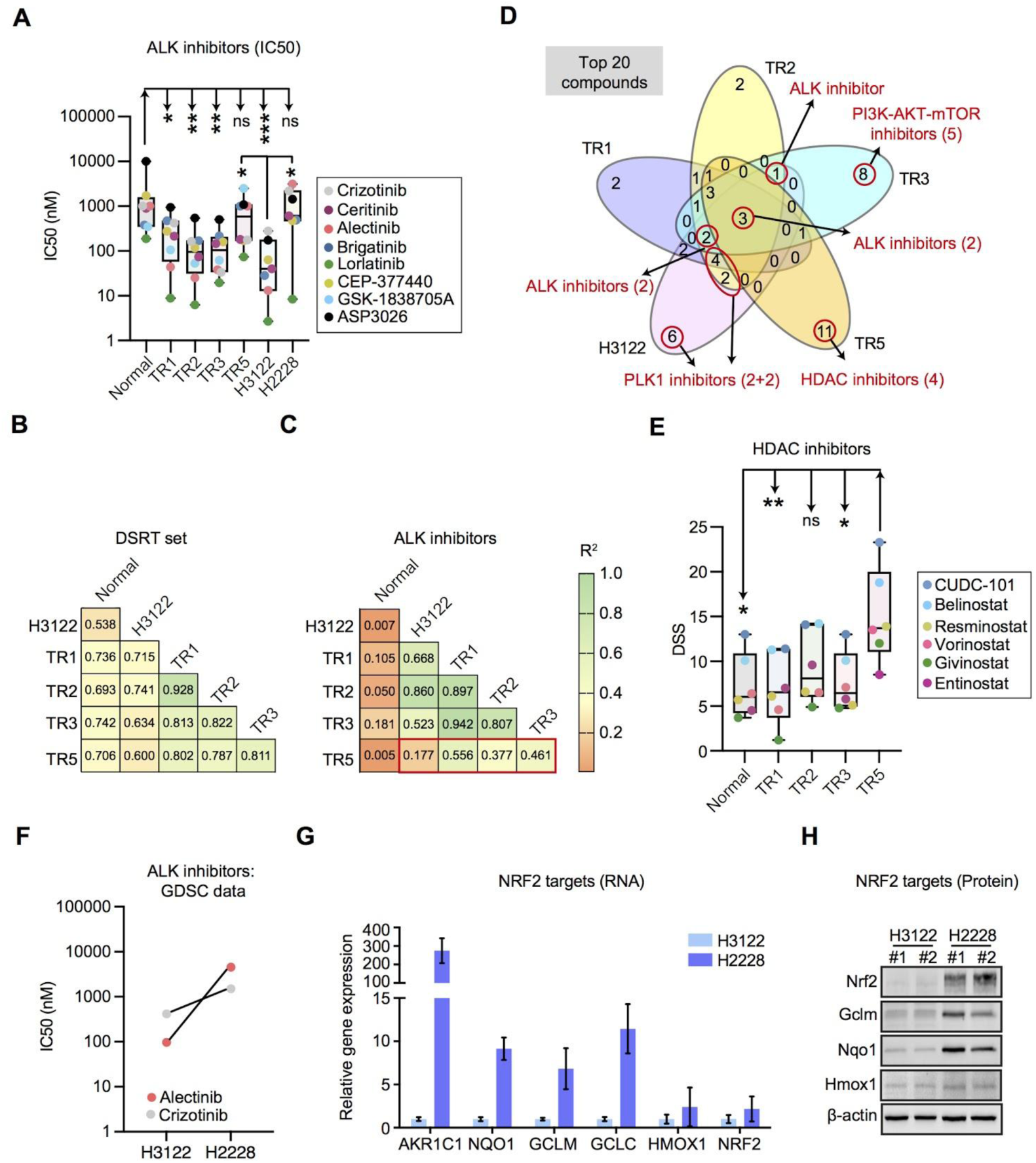
*ALK*-rearranged lung cancer cells exhibit a range of ALKi sensitivities. (A) The IC50 values of ALKi for indicated cell types. Each dot represents an individual drug and dots are color-coded based on drug ID. (B, C) Correlation analyses showing Pearson correlation coefficient values derived from DSS comparisons between two cell samples: comparisons were either made for (B) entire drug collection (n=527) or (C) only for drugs targeting ALK (n=8). (D) Comparisons of 20 compounds with highest selective DSSs for a particular cell type. (E) The DSSs of compounds targeting HDAC across different cell types. Each dot represents an individual drug and dots are color-coded based on drug ID. (F) The IC50 values of alectinib and crizotinib for H3122 and H2228 cells in the Genomics of Drug Sensitivity in Cancer (GDSC1) database. (G) Relative expression of Nrf2 and Nrf2 target genes measured by RT-qPCR, (H) Immunoblots of H3122 and H2228 cells probed with the indicated antibodies. Error bars represent minimum and maximum values or ± SEM. Student’s *t* test *p* values are * < 0.05, ** < 0.01, *** < 0.001.

**Figure S3.**
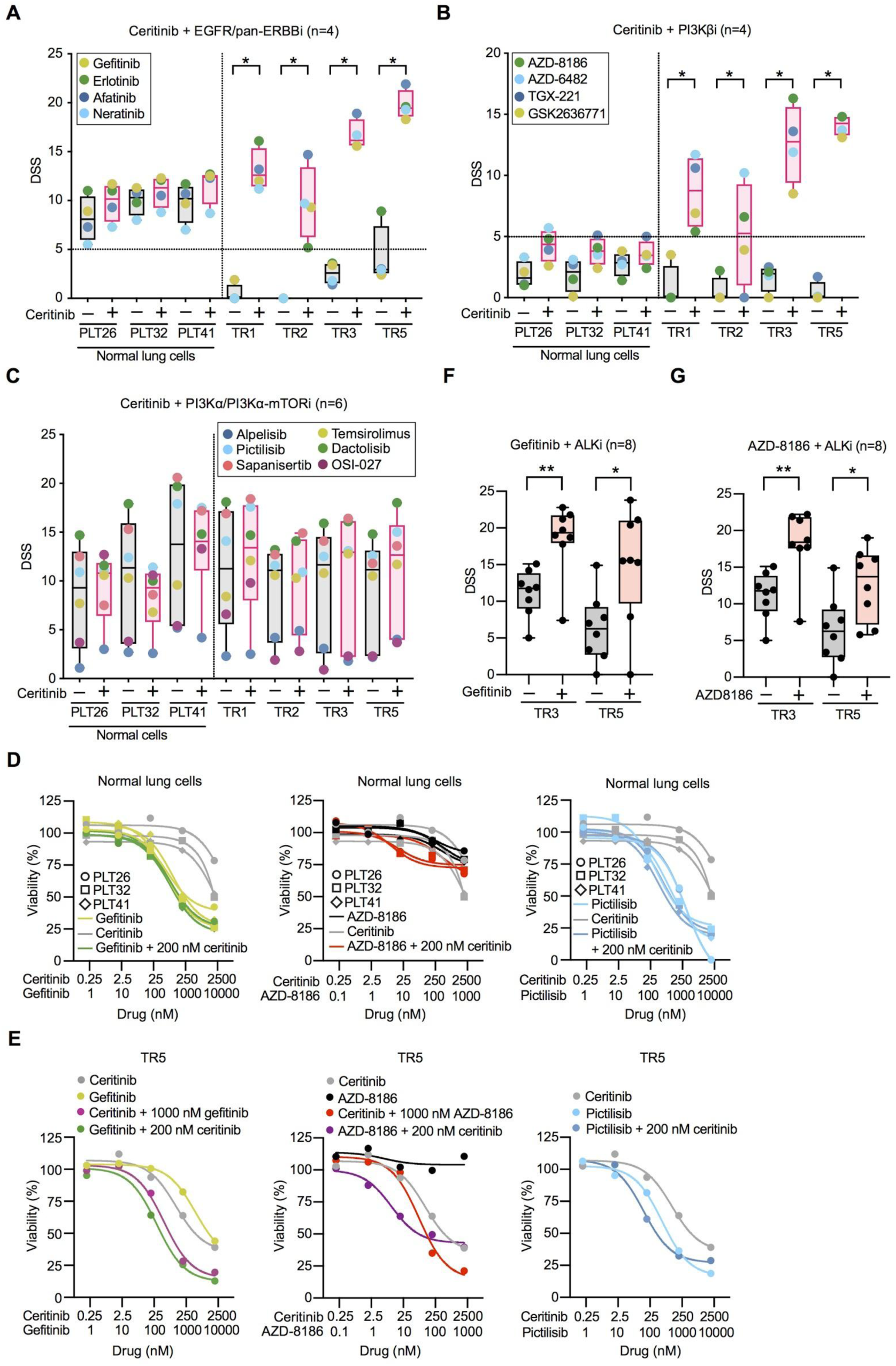
The effect of ALK and PI3Kβ inhibition is specific for the tumor cells. Normal lung or tumor-derived cells were treated with combination of (A) 200nM ceritinib + EGFRi or (B) 200 nM ceritinib + PI3Kβi or (C) 200 nM ceritinib + PI3Kα inhibitor; each dot in box chart represent the DSSs of single agents or combinations. Dose-response curves of ceritinib, gefitinib, AZD-8186, pictilisib or their combinations in (D) normal lung or (E) TR5 cells. The DSSs of ALKi as a single agent or in combination with (F) 1000 nM gefitinib or (G) 500 nM AZD-8186. Error bars represent minimum and maximum values. Student’s *t* test *p* values are * < 0.05, ** p < 0.01, *** p < 0.001.

**Figure S4.**
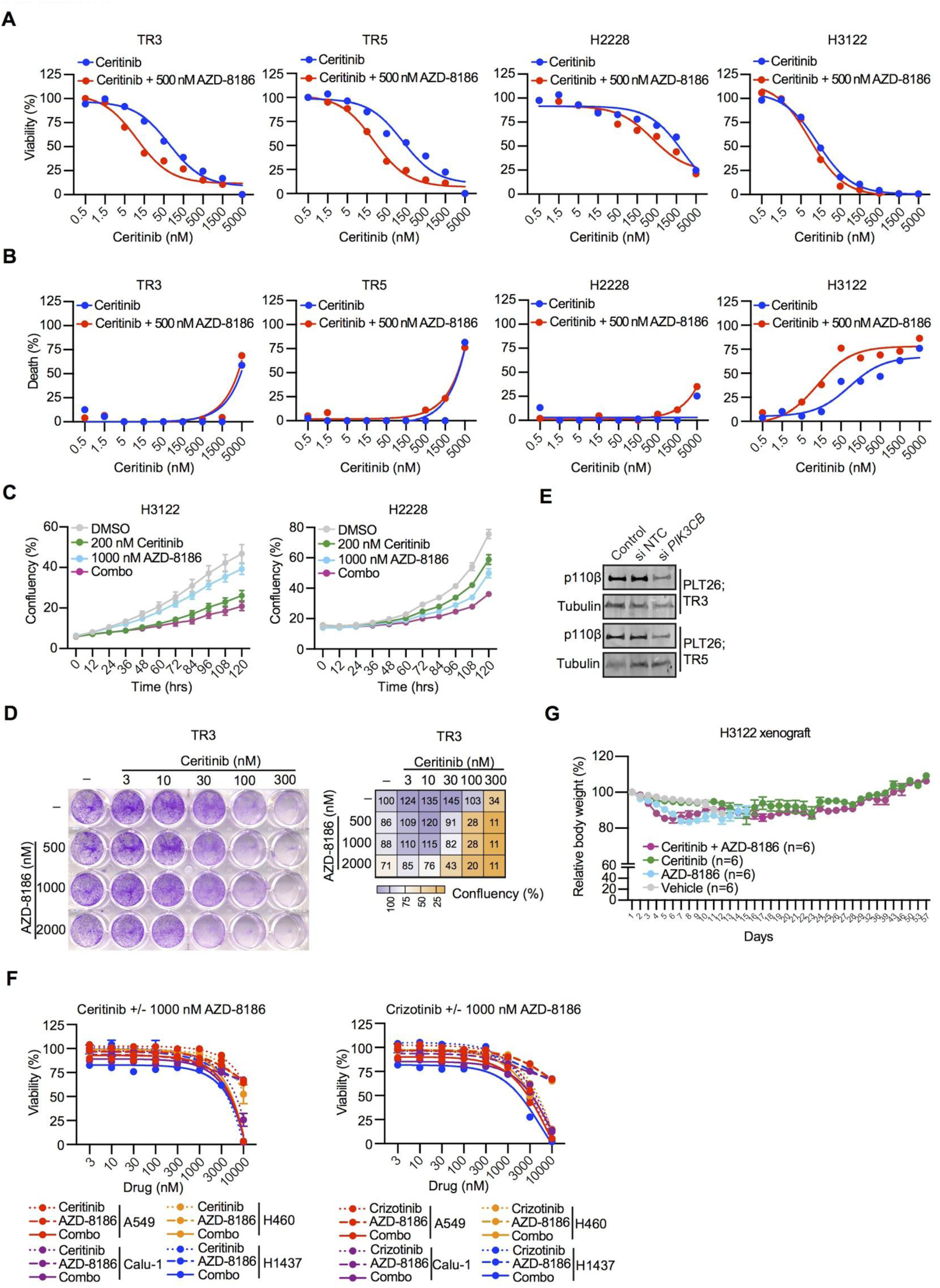
The PI3Kβi AZD-8186 increases ceritinib efficacy in *ALK*-rearranged lung cancer. Dose-response curves illustrating the effect of single agent ceritinib or combination of ceritinib plus 500 nM AZD-8186 on (A) cell viability or (B) cell death in TR3, TR5, H3122, and H2228 cells. (C) TR3 and H3122 cells were treated with vehicle (DMSO), 200 or 10 nM ceritinib, 1000 nM AZD-8186 or the combination of ceritinib plus AZD-8186; cell confluency was measured every 12 h for five days using an Incucyte live cell imaging system. (D) Representative images (left) of clonogenicity assays of TR3 cells treated for 15 days with five different concentrations of ceritinib, three different concentrations of AZD-8186, or their combination. Quantification (right) of colony formation assays. (E) Immunoblots of TR3 and TR5 cells transfected with NTC or *PIK3CB* siRNA and probed with the indicated antibodies. (F) *ALK* wildtype and *KRAS* mutant lung cancer cells (n=4) were treated with ceritinib, crizotinib, AZD-8186, or the combination of ceritinib or crizotinib plus 1000 nM AZD-8186. (G) Body weights of mice (monitored up to 57 days after treatment initiation) bearing subcutaneous H3122 xenografts that were administered with vehicle control, 25 mg/kg/day ceritinib, 2 × 25 mg/kg/day AZD-8186, or the combination of ceritinib and AZD-8186, for 21 days. Error bars represent ± SEM.

**Figure S5.**
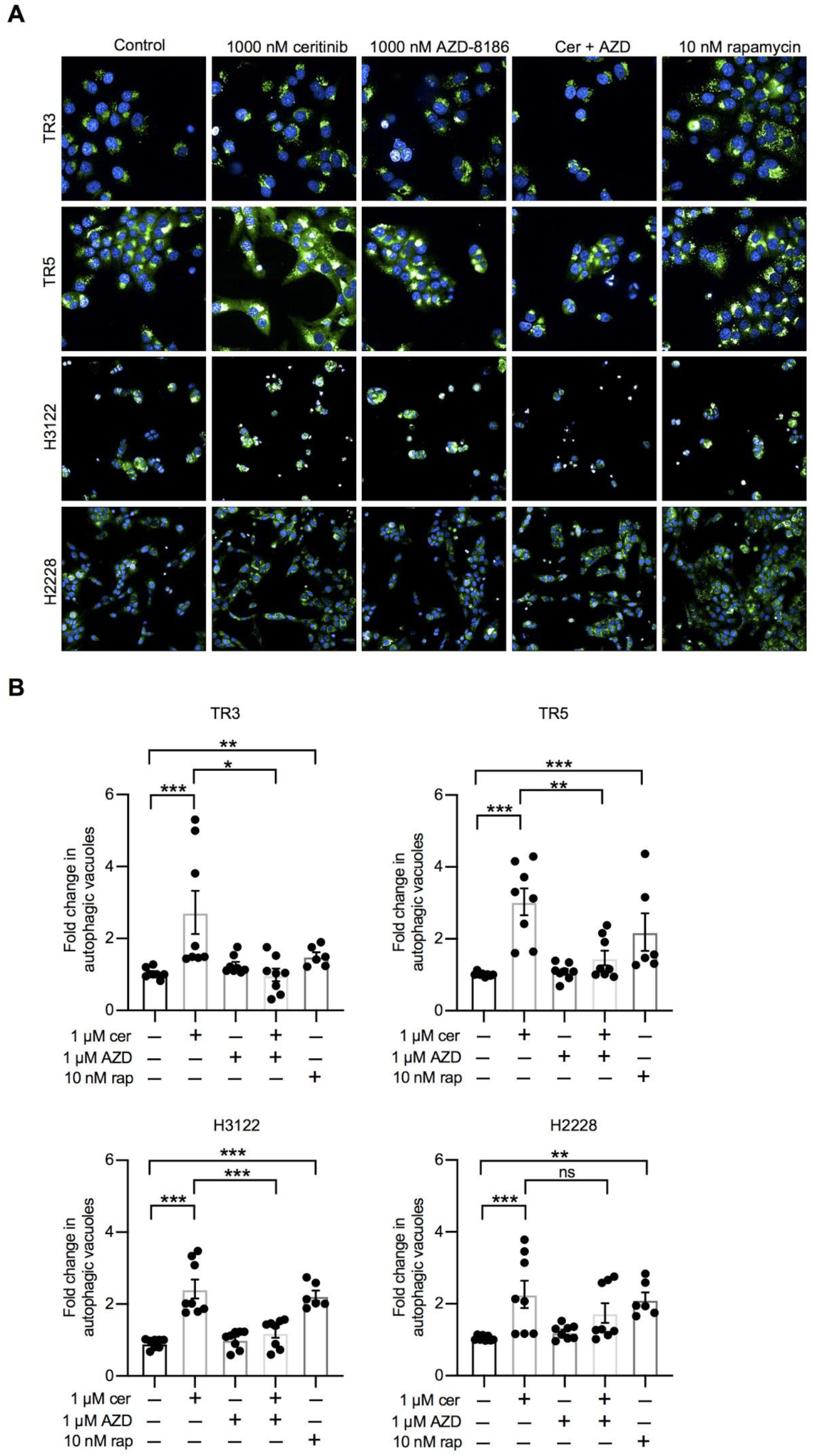
ALK inhibition leads to autophagy. (A) Representative images of TR5 and TR3 cells treated with vehicle control, 1000 nM ceritinib (cer), 1000 nM AZD-8186 (AZD), or their combination for 24 h. Before imaging, cells were stained with the Cyto-ID Green Detection Reagent to visualize autophagic vacuoles. Cells treated with 10 nM rapamycin (rap) served as a positive control. (B) Bar graph representing fold changes in the number of autophagic vacuoles in drug-treated cells relative to vehicle control. Error bars represent ± SEM. Student’s *t* test *p* values are * < 0.05, ** p <0.01, *** p <0.001.

**Figure S6.**
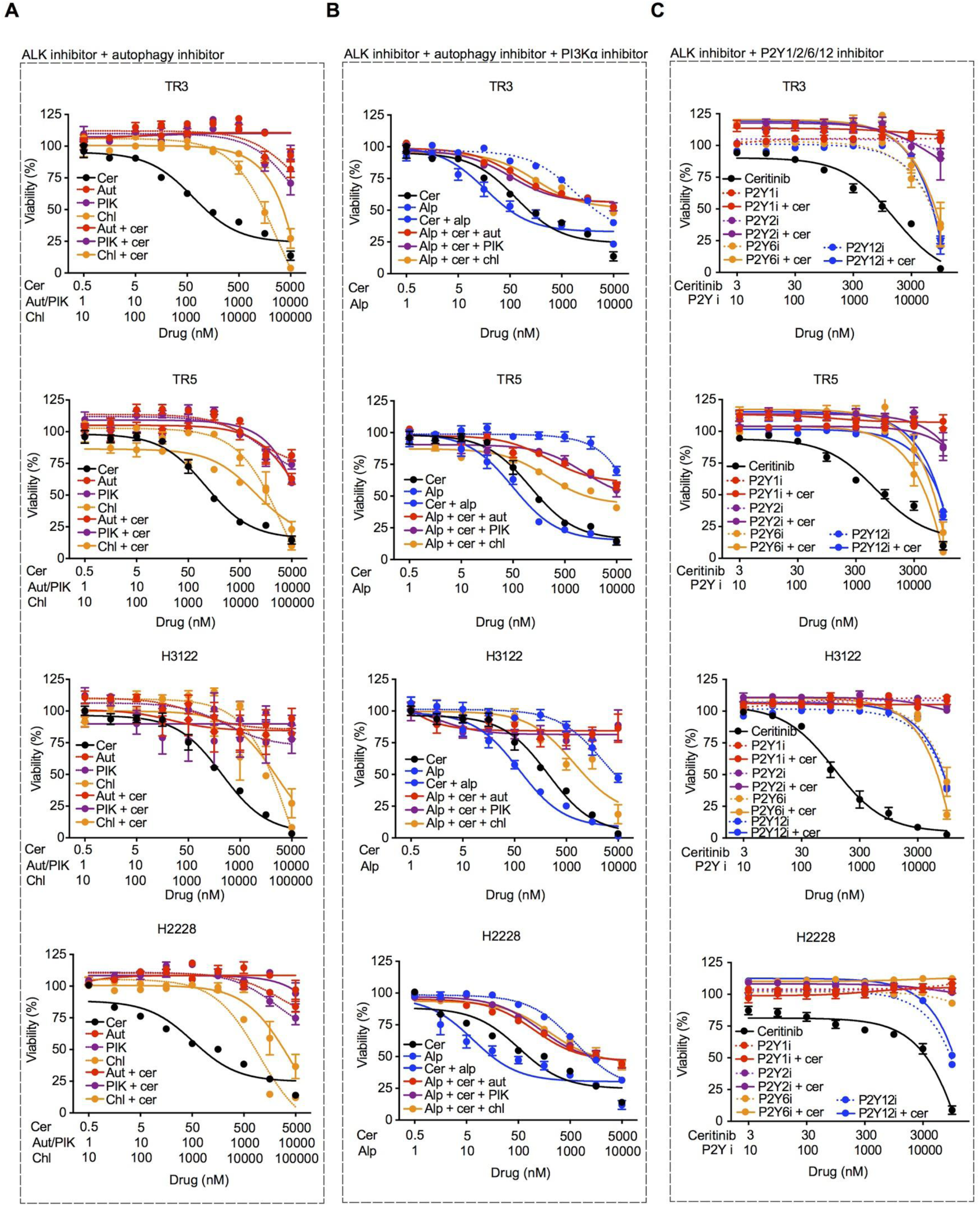
Inhibition of autophagy or P2Y receptors does not improve the response of ceritinib. (A) Dose-response curves of TR3, H2228 and H3122 cells treated with (A) ceritinib (cer), autophinib (aut; Vps34i), PIK-III (PIK; Vps34i), chloroquine (chl; autophagy inhibitor) or combinations of ceritinib plus 300 nM autophinib/PIK-III/chloroquine, (B) ceritinib, alpelisib (alp) or alpelisib in combination with 300 nm ceritinib or alpelisib in combination with 300 nm ceritinib plus 300 nM autophinib or 300 nM PIK-III or 10 µM chloroquine and (C) ceritinib, MRS2179 (P2Y1i), AR-C118925XX (P2Y2i), MRS2578 (P2Y6i), ticagrelor (tica; P2Y12i), or combination of P2Y inhibitors plus 200 nM ceritinib. Error bars represent ± SEM.

**Figure S7.**
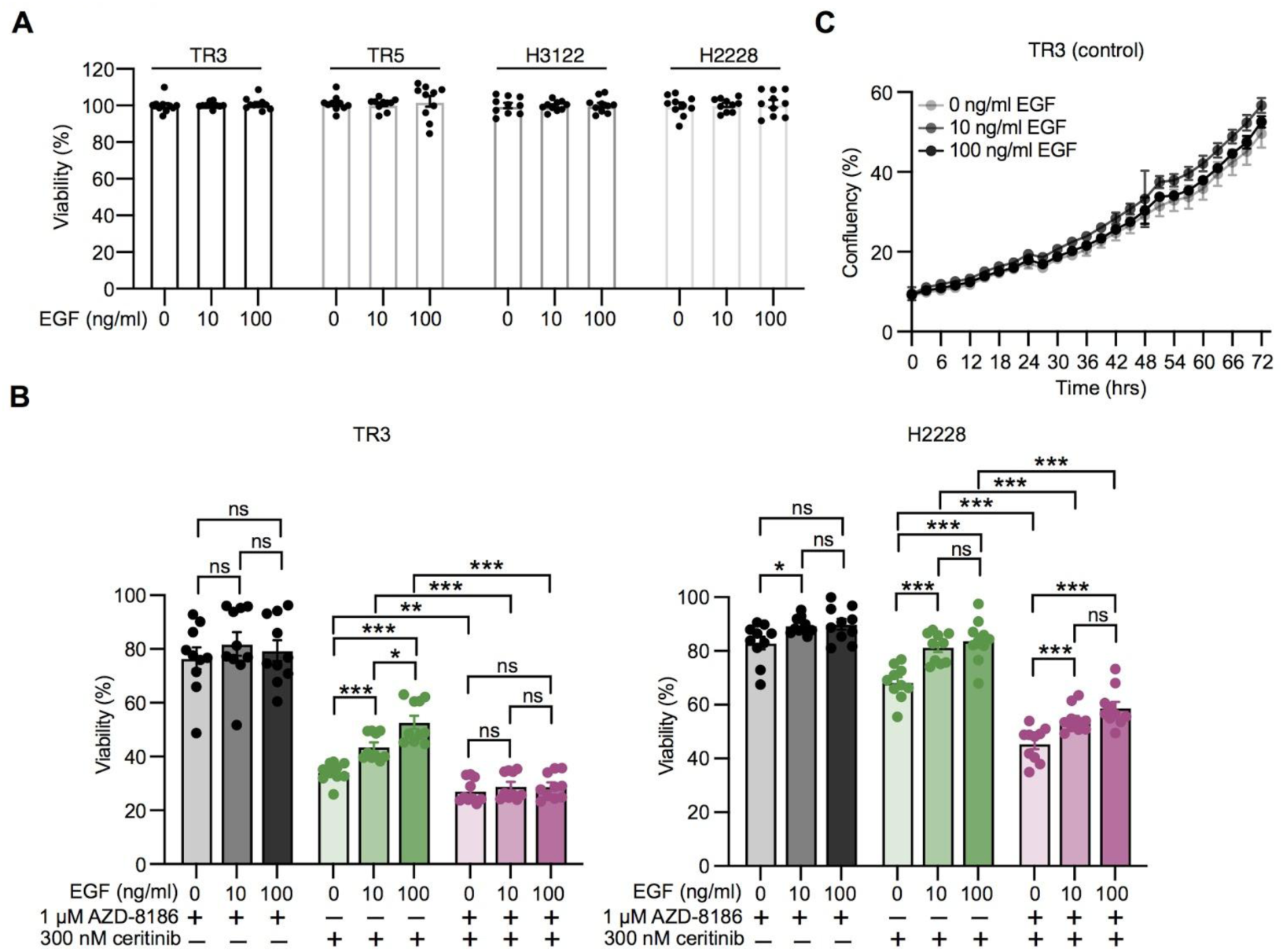
Combined inhibition of ALK and PI3Kβ overcomes EGFR mediated resistance in *ALK*-rearranged lung cancer cells. (A) Cell viabilities of TR3, TR5, H3122 and H2228 DMSO control cells are treated with different doses of EGF (0, 10, and 100 ng/ml). (B) TR3 and H2228 cells were co-treated with different doses of EGF (0, 10, and 100 ng/ml) and ceritinib or AZD-8186 or their combination for 72 h. Percentage viabilities of drug-treated cells were normalized to cells co-treated with different doses of EGF (0, 10, and 100 ng/ml) or DMSO. (C) TR3 control (DMSO) cells were treated with different doses of EGF (0, 10, and 100 ng/ml); cell confluency was measured every 3 h for 72 h using the Incucyte live cell imaging system.

**Table S1.** Details of primary antibodies used in immunohistochemistry and western blotting analyses

| **Immunohistochemistry** |  |  |  |
| --- | --- | --- | --- |
| **Antibody** | **Company** | **Catalog no. (clone)** | **Antigen retrieval** |
| NKX2-1 | Abcam | ab133638 (EPR8190-6) | 10 mM sodium citrate (pH 6.0) |
| LKB1 | Cell Signaling Technology | 13031 (D60C5F10) | 10 mM sodium citrate (pH 6.0) |
| Ki-67 | Thermo Fisher Scientific | RM-9106-S0 (SP6) | 10 mM sodium citrate (pH 6.0) |
| E-cadherin | Cell Signaling Technology | 3195 (24E10) | 10 mM sodium citrate (pH 6.0) |
| Pan-Cytokeratin (pan-CK) | Ventana | 760-2595 (AE1/AE3/PCK 26) | ULTRA Cell Conditioning Solution (CC1) from Ventana (pH 8.6) |
| Cytokeratin 18 (CK 18) | Dako | M7010 (DC-10) | ULTRA Cell Conditioning Solution (CC1) from Ventana (pH 8.6) |
| Vimentin | Abcam | ab92547 (EPR3776) | 10 mM sodium citrate (pH 6.0) |
| CD31 | Abcam | ab28364 (Rabbit polyclonal) | Tris-EDTA (pH 9.0) |
| Collagen IV | Roche | 760-2632 (CIV22) | Cell Conditioning Solution (CC2) from Ventana (pH 8.6) |
| **Immunoblotting** |  |  |  |
| **Antibody** | **Company** | **Catalog no. (clone)** | **Dilution/concentration** |
| α-Tubulin | Cell Signaling Technology | 2125 | 1:1000 |
| pAKT (Ser473) | Cell Signaling Technology | 4058 | 1:1000 |
| AKT | Cell Signaling Technology | 2920 | 1:1000 |
| pERK (Thr202/Tyr204) | Cell Signaling Technology | 4370 | 1:1000 |
| ERK | Cell Signaling Technology | 9107 | 1:1000 |
| pALK (Tyr1604) | Cell Signaling Technology | 3341 | 1:1000 |
| ALK | Cell Signaling Technology | 3633 | 1:1000 |
| pEGFR (Tyr1068) | Cell Signaling Technology | 3777 | 1:1000 |
| EGFR | Cell Signaling Technology | 4267 | 1:1000 |
| PI3Kβ | Cell Signaling Technology | 3011 | 1:1000 |
| Cleaved PARP (Asp214) | Cell Signaling Technology | 9541 | 1:1000 |
| **EGFR co-immunoprecipitation** |  |  |  |
| **Antibody** | **Company** | **Catalog no. (clone)** | **Amount** |
| EGFR | Santa Cruz Biotechnology | sc-120 | 1 μg |

